# Cell vulnerability within the sublaterodorsal tegmental nucleus underlies REM sleep behaviour disorder in prodromal α-synucleinopathy

**DOI:** 10.64898/2026.04.28.719687

**Authors:** Russell Luke, Han Soo Yoo, Dillon McKenna, Andrea Bevan, Casey Durso, Brittany J. Dugan, Yuling Liang, Bardia Amanirad, Jefferson A. Steltz, Bin Zhang, Allison Yearwood, Jonah Lourie, Phil Hyu Lee, Jimmy J. Fraigne, John Peever, Kelvin C. Luk

## Abstract

REM sleep behaviour disorder (RBD) is a prodromal manifestation of α-synucleinopathies such as Parkinson’s disease. Evidence suggests that degeneration of REM sleep regulating neurons in the sublaterodorsal tegmental nucleus (SLD) could give rise to RBD, yet the specific cellular populations involved and their contribution to synucleinopathy progression remain unclear. Here, we investigated the role of defined SLD cell types in RBD pathogenesis. Using viral vector and fibril-based models of α-synucleinopathy, we show that α-synuclein pathology in SLD neurons triggers RBD in mice. Notably, glutamatergic SLD neurons are selectively vulnerable to synucleinopathy and the loss of these cells correlates with RBD severity. Propagation of synucleinopathy from the SLD to midbrain and forebrain structures leads to the emergence of neurological deficits associated with Parkinson’s disease. These findings establish that SLD neurons are critical substrates for RBD and provide insight into the cellular mechanisms at play in the early stages of synucleinopathies.

## INTRODUCTION

Parkinson’s disease (PD) is a neurodegenerative disorder characterized by bradykinesia, rigidity and tremors, and biologically defined by the intracellular aggregation of α-synuclein in neurons and neurites^1,2^. The discovery that the motor symptoms of PD arise from α-synuclein mediated degeneration of dopaminergic midbrain neurons has advanced our understanding of the disease. PD is now widely recognized as a multisystem synucleinopathy with a lengthy prodromal phase, during which non-motor symptoms precede the onset of classical motor deficits. For example, changes in olfactory and gut/urinary function often precede the onset of classic PD by a decade or more^3–5^.

Identifying how different cell types are affected by synucleinopathy during the early stages of PD is central to understanding the basis of its disease etiology and progression. Among these prodromal symptoms, REM sleep behaviour disorder (RBD) has emerged as the strongest clinical predictor of PD^6–8^. RBD is characterized by the loss of normal muscle paralysis/atonia that accompanies REM sleep, resulting in excessive movements during REM sleep that can lead to serious patient or bed partner injuries^9,10^. Roughly half of PD patients present with RBD^11–13^, while 80-90% of people with RBD develop PD or a related α-synucleinopathy (e.g., dementia with Lewy bodies) within 5-15 years of diagnosis. Despite this strong association, it remains unclear why RBD patients develop α-synucleinopathy. A longstanding hypothesis that accounts for the link between RBD and PD is that they both share the common mechanism, whereby α-synuclein (αSyn) pathology initially targets brainstem circuits that generate REM sleep atonia, before spreading rostrally into the midbrain dopamine circuits associated with PD^14,15^.

Experimental and clinical studies indicate that the sublatedorosal tegemental nucleus (SLD) or Subcoeruleus nucleus (SubC) in humans, generates natural REM sleep atonia. Lesions of the SLD impair REM sleep atonia and induce RBD behaviours in both animals and humans^16–21^. In addition, imaging and post-mortem studies reveal damage or α-synuclein pathology in this region is associated with RBD in PD patients^7,24^. These findings establish the SLD as an anatomical substrate linking REM sleep dysfunction with early synucleinopathic disease, yet they do not identify which neuronal populations within the SLD are most vulnerable to pathology and those that are responsible for RBD symptomatology. The specific cellular substrates within the SLD that confer vulnerability to synucleinopathy, and how pathology in these cells contributes to progression from RBD to PD remains elusive.

Here, we sought to identify the SLD cells that are vulnerable to synucleinopathy and determine how they contribute to RBD pathogenesis and disease progression. To do this, we used a viral vector-mediated approach to selectively overexpress αSyn in SLD glutamate neurons and then we injected αSyn PFFs into the SLD of wild-type mice to initiate the spread of αSyn pathology from this region and evaluated resulting pathological and behavioural outcomes. We found that SLD glutamate neurons exhibited pronounced vulnerability to αSyn pathology which induced robust RBD motor behaviours. The progressive spread of pathology from these cells to other structures, including the substantia nigra, pedunculopontine nucleus, lateral hypothalamus and ventrolateral preoptic nucleus, led to the emergence of Parkinsonian motor deficits and additional sleep disturbances. Together, these findings establish a mechanistic framework by which αSyn pathology in defined SLD cell types can initiate the progression of synucleinopathy from prodromal to advanced disease stage.

## RESULTS

### αSyn pathology in the SubC in Lewy body disease (LBD) patients with RBD

To confirm the association between subcoeruleal αSyn pathology and RBD, we compared the pathological burden in the SubC (homologous to the rodent SLD) of healthy control individuals, LBD subjects without RBD, and those with RBD, obtained at autopsy. Immunostaining against αSyn phosphorylated at Ser129 (p-αSyn) revealed abundant αSyn pathology in the SubC of LBD patient samples compared to the control samples (Fig. 1a-c), with p-αSyn-positive area in the SubC being higher in LBD patients with RBD than those without RBD, although this comparison did not reach statistical significance (Fig. 1d). These results suggest that the presence of pathological αSyn aggregates in the SubC is associated with the manifestation of RBD in humans.

**Fig. 1.**
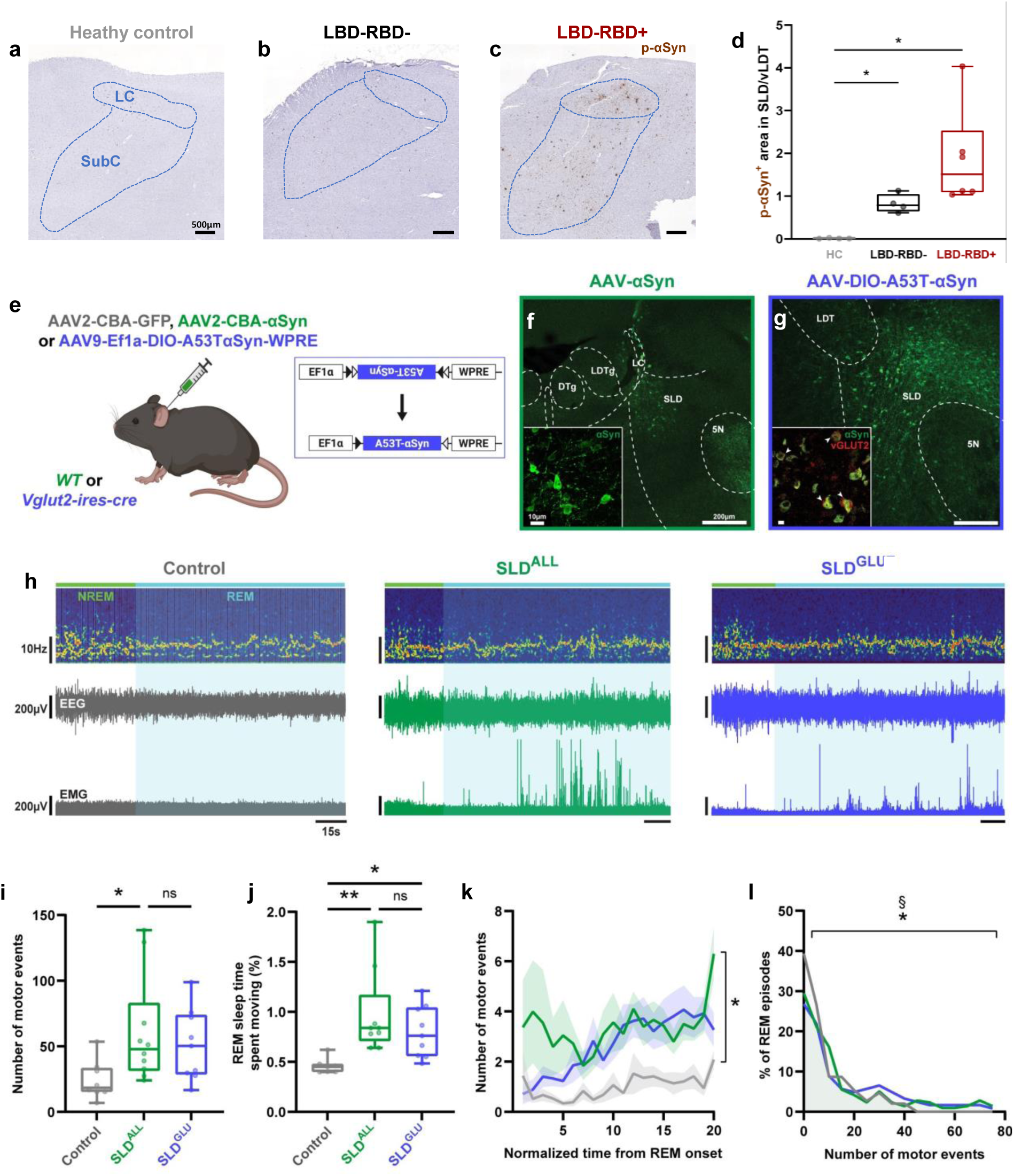
αSyn pathology in the SubC/SLD was associated with RBD in humans and mice. **(a-c**) Immunohistochemical staining for phosphorylated αSyn (p-αSyn, brown) in post-mortem brain samples from healthy elderly individuals (HC) (**a**), Lewy body disease patients without RBD diagnosis (LBD-RBD-) (**b**), and Lewy body disease patients with comorbid RBD (LBD-RBD+) (**c**). The pontine areas corresponding to the locus coeruleus (LC) and subcoeruleus (SubC) are outlined in blue. Note the robust staining of αSyn aggregates in the SubC of patients with RBD, compared to PD patients without RBD or healthy controls. (**d**) Quantitation of histological data reveals a gradient of pathology distributed among the PD patient population depending on RBD status, such that p-αSyn immunoreactivity was increased in the SubC of patients with RBD (red box) compared to healthy controls (grey box) (\**P* < 0.05, *n* = 4-6). RBD-negative LBD patients (black box) tended to develop αSyn aggregates to a lesser degree than RBD-positive cases but still had significantly more αSyn aggregates compared to healthy controls. (**e)** Overview of experimental strategy for pan-neuronal (green) and glutamatergic cell-specific (violet) αSyn overexpression in SLD neurons. (**f-g**) Representative images show immunofluorescent staining of αSyn (green) in SLD neurons **(f)** and selective expression of αSyn (white arrowheads) within vGLUT2-positive SLD neurons (red) at 8-10 weeks post-AAV injection **(g)**. (**h)** Example electrophysiological recordings from mice injected with AAV-GFP (grey), AAV-αSyn (green), or AAV-DIO-A53T-αSyn (violet) at 8-10 weeks post-injection. Note the heightened EMG activity during REM sleep (cyan) in both groups of αSyn-transduced mice. (**i**-**l)** EMG analyses reveal that pan-neuronal and glutamatergic cell-specific αSyn overexpression in the SLD injection increased REM sleep motor activity, as indicated by an increase in the number of motor events during REM sleep (**i**) (Control vs. SLD^ALL^: \**P* < 0.05; Control vs. SLD^GLU^ *P* = 0.0712, *n* = 8-10) and a greater proportion of REM sleep time occupied by these movements (**j)** (Control vs. SLD^ALL^: **P < 0.01, Control vs. SLD^GLU^: *P <0.05, n = 8-10) **P < 0.01,\**P* < 0.05, *n* = 8-10). REM sleep bouts were standardized by duration, and motor events were quantified across each bout. **(k)** Group data showing that the pattern of movements within individual REM sleep episodes is altered following pan-neuronal (green) and cell-specific (violet) αSyn overexpression, such that both experimental groups displayed more movements across REM sleep episodes compared to GFP controls (grey line) (Main treatment effect: \**P* < 0.05, *n* = 8-10). **(l)** Both groups of αSyn-transduced mice displayed a shift in the frequency distribution of REM sleep bouts towards bouts with more movements compared to REM sleep bouts from control mice (Control vs. SLD^ALL^: \**P* < 0.05, Control vs. SLD^GLU^: §*P* < 0.05, *n* = 146-262 bouts).

### Development of RBD phenotype in mice with αSyn overexpression in the SLD neurons

We next investigated whether elevated αSyn levels in the SLD triggers an RBD phenotype *in vivo* by injecting wild-type mice with an adeno-associated virus (AAV) encoding wild-type human ɑSyn into the SLD (Fig. 1e). Electrophysiological studies were conducted at 8-10 weeks post-AAV injection to quantify tonic and phasic components of muscle activity during REM sleep. Mice overexpressing ɑSyn in SLD neurons exhibited a marked increase in REM sleep motor activity, as reflected by increases in the number and total duration of phasic motor events compared to control animals which exhibited minimal muscle activity during REM sleep (Fig. 1h-j). Correspondingly, within-episode analysis of motor activity further revealed that motor events were more frequent across individual REM sleep episodes, and that the distribution of REM sleep episodes shifted towards episodes that contained more movements in ɑSyn-transduced mice (Fig. 1k, l). Despite heightened motor activity during REM sleep, neither EEG activity nor the timing of REM sleep itself were affected by ɑSyn overexpression (Supplementary Fig. 1). Together, these findings underscore a selective function of SLD neurons in suppressing motor activity during REM sleep and establish that αSyn overexpression in these neurons is sufficient to elicit RBD behaviours in mice.

Glutamatergic SLD neurons have been identified as a principal cellular substrate for REM sleep muscle atonia^25–27^. To address the role of this neuronal subpopulation in driving the observed RBD-like phenotype, we selectively overexpressed αSyn in glutamatergic SLD neurons by injecting AAV-DIO-A53T-ɑSyn into the SLD of Vglut2-ires-cre mice (Fig. 1e,g). Targeted expression of ɑSyn in glutamatergic SLD neurons recapitulated RBD-like behaviours observed following pan-neuronal overexpression, resulting in increased REM sleep motor activity compared to controls. Specifically, cell-specific αSyn overexpression increased the total duration of phasic motor events during REM sleep and caused a trend towards a higher number of these movements (Fig. 1j,k). The temporal pattern of movements within REM sleep bouts and the distribution of REM sleep bouts were likewise consistent with pan-neuronal experiments, with cell-specific ɑSyn overexpression leading to elevated motor activity throughout individual REM sleep bouts and a greater proportion of bouts with higher phasic motor output. Neither the amount of REM sleep nor its EEG characteristics were affected by αSyn overexpression in glutamatergic SLD neurons (Supplementary Fig. 1), indicating a selective disruption of REM sleep atonia. These findings demonstrate that ɑSyn overexpression in glutamatergic SLD neurons is sufficient to induce RBD-like symptoms in mice, establishing this subpopulation as a major cellular substrate underlying RBD symptoms.

### Fibril-based induction of αSyn pathology in the SLD and its propagation throughout the CNS connectome

Human post-mortem studies indicate that the SubC is affected by Lewy pathology at the onset of iRBD^19–21^, whereas individuals with iRBD show positivity on CSF or serum ɑSyn seed amplification assays^28,29^. This suggests that the expansion of pathology beyond the SLD may underlie progression from iRBD into advanced motor and non-motor phenotypes of LBD. To determine whether synucleinopathy spreads to surrounding and distal neuronal populations over time, we stereotaxically injected recombinant αSyn PFFs into the SLD of wildtype mice to initiate propagative synucleinopathy from this region (Fig. 2a). Separate cohorts of mice were analyzed by p-ɑSyn immunohistology at 1, 3, 6 or 12 months following a single unilateral administration of PFFs to map the spatiotemporal distribution of ɑSyn pathology across the CNS connectome (Fig. 2b-d). Beginning at 1 mpi, pathology was detectable in the SLD injection site and neighboring ventral laterodorsal tegmental nucleus (vLDT) (Fig. 2c,d), as well as in the ventral medulla, parabrachial nuclei, pedunculopontine nuclei, and periaqueductal grey (Fig. 2d,e). By 3 mpi, pathology was present in additional caudal (e.g., dorsal motor nucleus of vagus and spinal cord) and rostral (e.g., midbrain reticular nucleus, amygdala, and bed nuclei of stria terminalis) regions, indicating further transneuronal propagation of pathology. At 6 mpi, neuronal αSyn inclusions were found in the substantia nigra pars compacta and ventral tegmental area. Hypothalamic structures, including the anterior and lateral hypothalamic areas and ventrolateral preoptic nucleus, parafascicular nucleus of the thalamus, nucleus accumbens, and infralimbic area, were also affected. By 12 mpi, pathology was further detectable in the tuberomammillary nucleus and anterior cingulate area. Interestingly, pathological burden in the SLD, initially increased over time before gradual dissipating, consistent with neuron loss as previously described for other CNS regions in this experimental paradigm^30,31^.

**Fig. 2.**
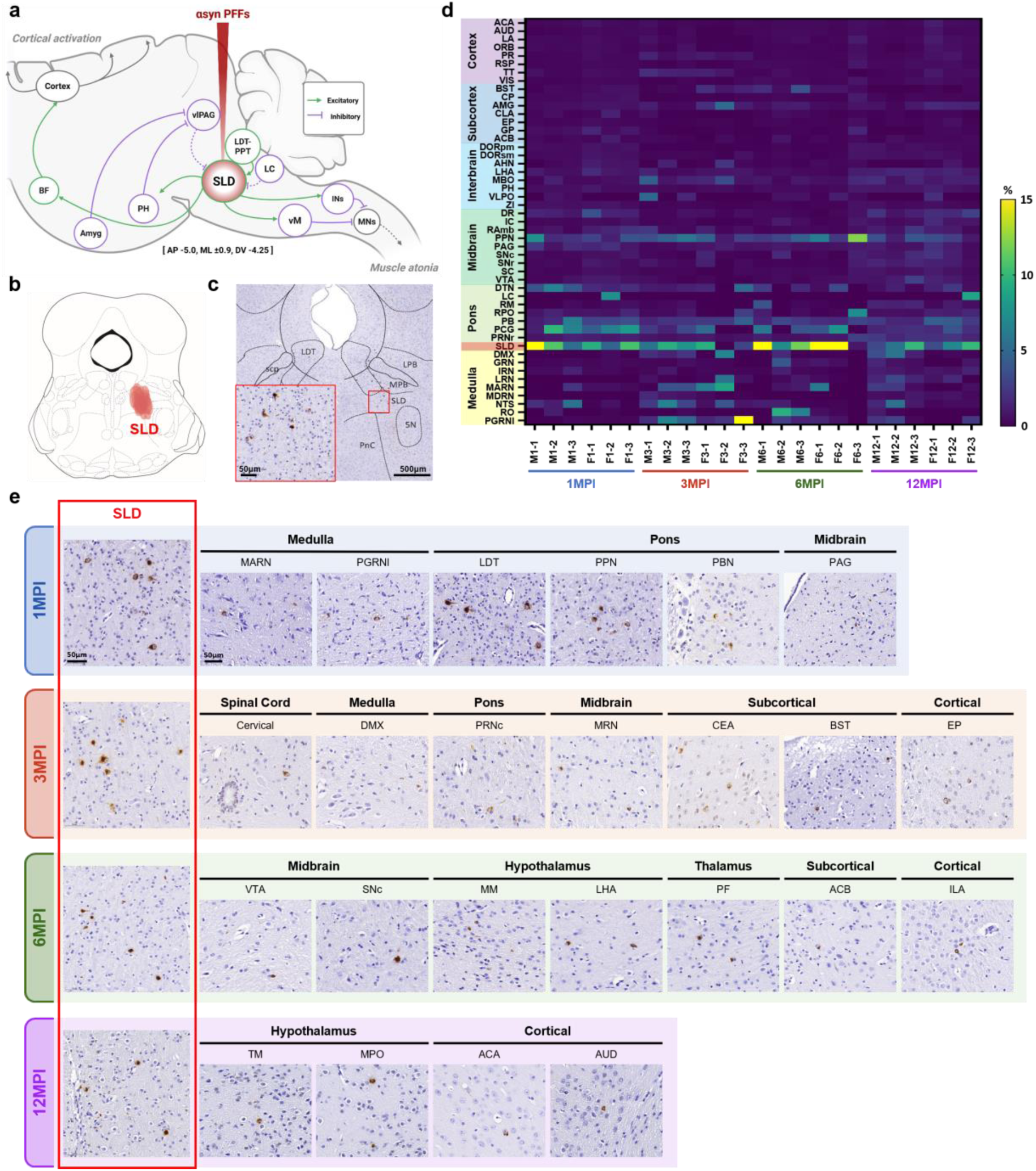
αSyn PFF inoculation into the SLD seeded the propagation of αSyn pathology throughout the CNS. **(a**) Schematic shows the hypothesized neural circuit responsible for generating cortical activation and muscle atonia during REM sleep. αSyn PFFs were injected into the SLD the main neural substrate of REM sleep. Green lines indicate excitatory pathways, and purple lines indicate inhibitory pathways. (**b**) 500nL of preformed-fibrils composed of wild-type mouse αSyn into the right SLD of wild-type mice. Red areas shows the stereotaxic injection sites for 15 mice. **c**, Representative coronal brain section showing the spatial distribution of p-αSyn aggregates (brown staining) in the pons at 1 mpi, with high magnification inlay. (**d**) Heatmap shows automatic quantification of pathological αSyn burden across various brain regions. Pathological burden was defined as the percentage of p-αSyn positive area divided by total areas of each region. Brain regions were ordered on the y-axis from medulla (bottom) to cerebral cortex (top). Results for a total of 24 mice are shown along the x-axis from 1 mpi (left) to 12 mpi (right). Red box denotes SLD as the injection site. (**e**) Representative images of p-αSyn staining in PFF-injected mice from 1 to 12 mpi. Representative images of the SLD were placed on the left side of each time point to show the progression of pathological burden in the SLD/vLDT longitudinally. Abbreviations = ACA, anterior cingulate area; ACB, nucleus accumbens; Ach, acetylcholine; AHA, anterior hypothalamic area; AMY, amygdala; AON, anterior olfactory nucleus; AP, anteroposterior; AUD, auditory area; BF, basal forebrain; BST, bed nuclei of the stria terminalis; CLA, claustrum, CNU, cerebral nuclei; CP, caudoputamen; CTX, cerebral cortex; DMX, dorsal motor nucleus of the vagus nerve; DORpm, polymodal association cortex related thalamus; DORsm, sensory-motor cortex related thalamus; DR, dorsal nucleus raphe; DTN, dorsal tegmental nucleus; DV, dorsoventral; EP, endopiriform nucleus; F, female; Glut, glutamate; Gly, glycine; GP, globus pallidus; GRN, gigantocellular reticular nucleus; IC, inferior colliculus; ILA, infralimbic area; ILM, intralaminar nuclei of the thalamus; INs, motor neurons in the spinal cord; IRN, intermediate reticular nucleus; LA, limbic area; LC, locus coeruleus; LDT, laterodorsal tegmental nucleus; LHA, lateral hypothalamic area; LPB, lateral parabrachial nucleus; LRN, lateral reticular nucleus; M, male; MARN, magnocellular reticular nucleus; MB, midbrain; MBO, mammillary body; MCH, melanin concentrating hormone; MDRN, medullary reticular nucleus; ML, mediolateral; MM, medial mammillary nucleus; MNs, motor neurons in the spinal cord; MPB, medial parabrachial nucleus; MPO, medial preoptic area; MRN, midbrain reticular nucleus; NE, norepinephrine; NTS, nucleus of the solitary tract; ORB, orbital area; PAG, periaqueductal gray; PB, parabrachial nucleus; PCG, pontine central gray; PF, parafascicular nucleus; PGRNl, lateral part of the paragigantocellular reticular nucleus; PH, posterior hypothalamic nucleus; PIR, piriform area; PnC, caudal part of the pontine reticular nucleus; PPN, pedunculopontine nucleus; PRNr, pontine reticular nucleus; RAmb, midbrain raphe nuclei; RM, raphe magnus; RO, nucleus raphe obscurus; RPO, nucleus raphe pontis; RSP, retrosplenial area; SC, superior colliculus; SCP, superior cerebellar peduncle; SLD/vLDT, sublaterodorsal tegmental nucleus/ventral part of laterodorsal tegmental nucleus; SNc, compact part of the substantia nigra; SNr, reticular part of the substantia nigra; TM, tuberomammillary nucleus; TT, taenia tecta; VIS, visual area; vlPAG, ventrolateral periaqueductal gray; VLPO, ventrolateral preoptic nucleus; vM, ventral medial medulla; VTA, ventral tegmental area; ZI, zona incerta; 5N, motor trigeminal nucleus.

### Synucleinopathy in the SLD leads to an RBD-like phenotype in mice

We next examined whether PFF-seeded synucleinopathy in the SLD and its subsequent propagation to other regions led to the development of a RBD phenotype in wildtype mice. RBD-like behaviours in mice were assessed using longitudinal EMG measurements and video footage collected between 3-9 mpi from animals that received either PFFs or a sham PBS injection into the SLD. PFF-injected mice developed an RBD-like phenotype as early as 3 mpi, the first time point examined. An independent cohort injected with αSyn PFFs bilaterally exhibited sustained increases in muscle activity during REM sleep, characterised by a higher frequency of phasic motor events (Fig. 3a,b), an extended duration of phasic motor movements in REM sleep bouts (Fig. 3c), and greater variability in muscle tone until 9 mpi (Fig. 3d). PBS-injected mice did not show changes in EMG activity over time (Supplementary Fig. 2a-c). Analysis of the distribution of phasic motor events across and within individual REM sleep bouts revealed that PFF-injection altered the distribution of REM sleep episodes, with PFF-treated mice exhibiting a higher proportion of REM sleep episodes having more motor events (Fig. 3e). This elevation in motor activity was maintained across the entire duration of REM sleep episodes (Fig. 3f,g). Mice that received unilateral PFF-injections developed a moderate and transient increase in EMG activity during REM sleep until 6 mpi (Supplementary Fig. 3). In both uni– and bilaterally inoculated mice, the changes in muscle activity were only observed during REM sleep but not present in NREM sleep or wakefulness (Supplementary Fig. 4), while no discernable differences in RBD-like behaviours were observed between male and female mice (Supplementary Fig. 5).

**Fig. 3.**
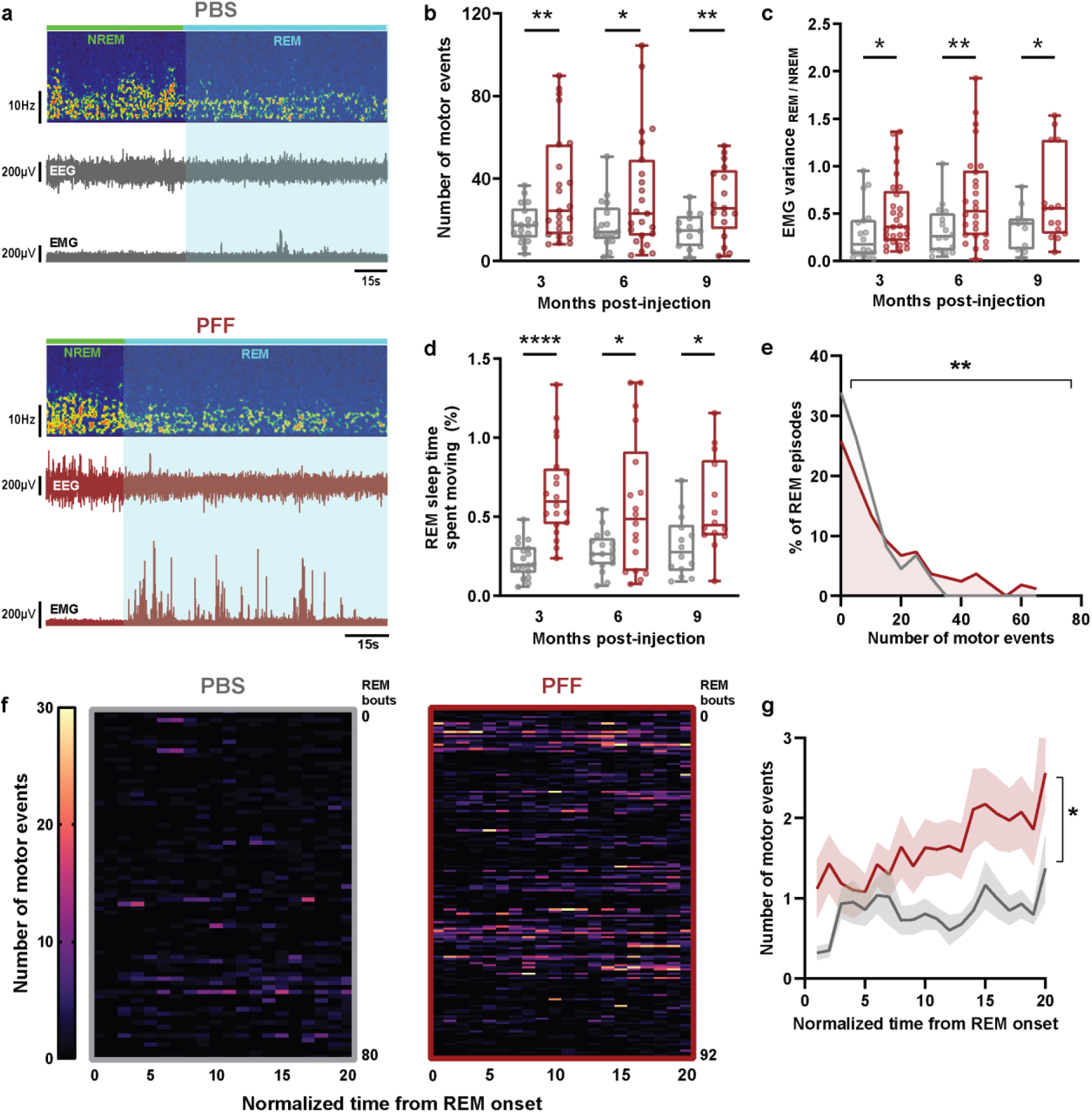
PFF injection into the SLD induced RBD-like behaviours characterised by heightened muscle activity during REM sleep. **(a**) Example electrophysiological recording from mouse injected with αsyn PFFs (red) in the SLD/vLDT at 9mpi and age-matched PBS control (grey). The PFF-injected mouse displayed exaggerated muscle activity during REM sleep (cyan). The preceding NREM sleep (green) is shown for comparison. (**b-c**) Quantification of EMG activity reveals that PFF injection increased the number of movements (**b**) (\**P* < 0.05, \*\**P* < 0.01, *n* = 18-22) and time spent moving (**c**) (\**P* < 0.05, \*\*\*\**P* < 0.0001, *n* = 18-22) during REM sleep between 3-9mpi. (**d**) PFF-injected mice displayed a higher ratio of EMG variance between REM and NREM sleep compared to age-matched controls (\**P* < 0.05, \*\**P* < 0.01, *n* = 18-22. (**e**) The frequency distribution of REM sleep episodes from PFF-injected mice was skewed towards episodes with more movements compared to the control population at 6mpi (\*\**P* < 0.01, *n* = 122-146 episodes). (**f)** Heatmaps show EMG activity across all REM sleep episodes from PBS– (left) and PFF-injected mice (right) at 6mpi. REM sleep bouts were standardized in length and the number of motor events were quantified along each REM episode. (**g**) Group data comparing the temporal profile of movements within individual REM sleep episodes from PFF-injected mice (red line) and age-matched PBS controls (grey line). The pattern of motor activity within REM sleep bouts from PFF-injected mice differed from that of controls at 6mpi, such that movements are more frequent across the duration of REM sleep bouts (\**P* < 0.05, *n* = 18-22).

### Changes in EEG frequency and sleep architecture in mice with SLD αSyn pathology

Although RBD is the most prominent sleep disturbance in early-stage PD, RBD patients also present with cortical EEG slowing in wakefulness and REM sleep^32,33^. Besides RBD, more generalized disruptions in sleep-wake architecture including insomnia and sleep fragmentation are also common at advanced stages of PD^34–37^. To evaluate the impact of PFF injection on cortical activation, we conducted spectral analysis of EEG activity across different vigilance states. Starting from 6 mpi, mice that received bilateral PFF inoculations exhibited a shift in EEG spectral composition toward lower frequencies, characterised by reduced theta power and correspondingly elevated delta power during NREM sleep, REM sleep and wakefulness (Fig. 4a-c). This widespread EEG slowing was further evidenced by a consistent reduction in the theta-to-delta power ratio across all vigilance states (Fig. 4d). Additionally, PFF injections led to increased wakefulness and a corresponding decrease in NREM and REM sleep duration starting from 6 mpi (Fig. 4e-g). This sleep loss appeared to be driven by prolonged bouts of wakefulness and reduced entry into sleep states (Fig. 4h).

**Fig. 4.**
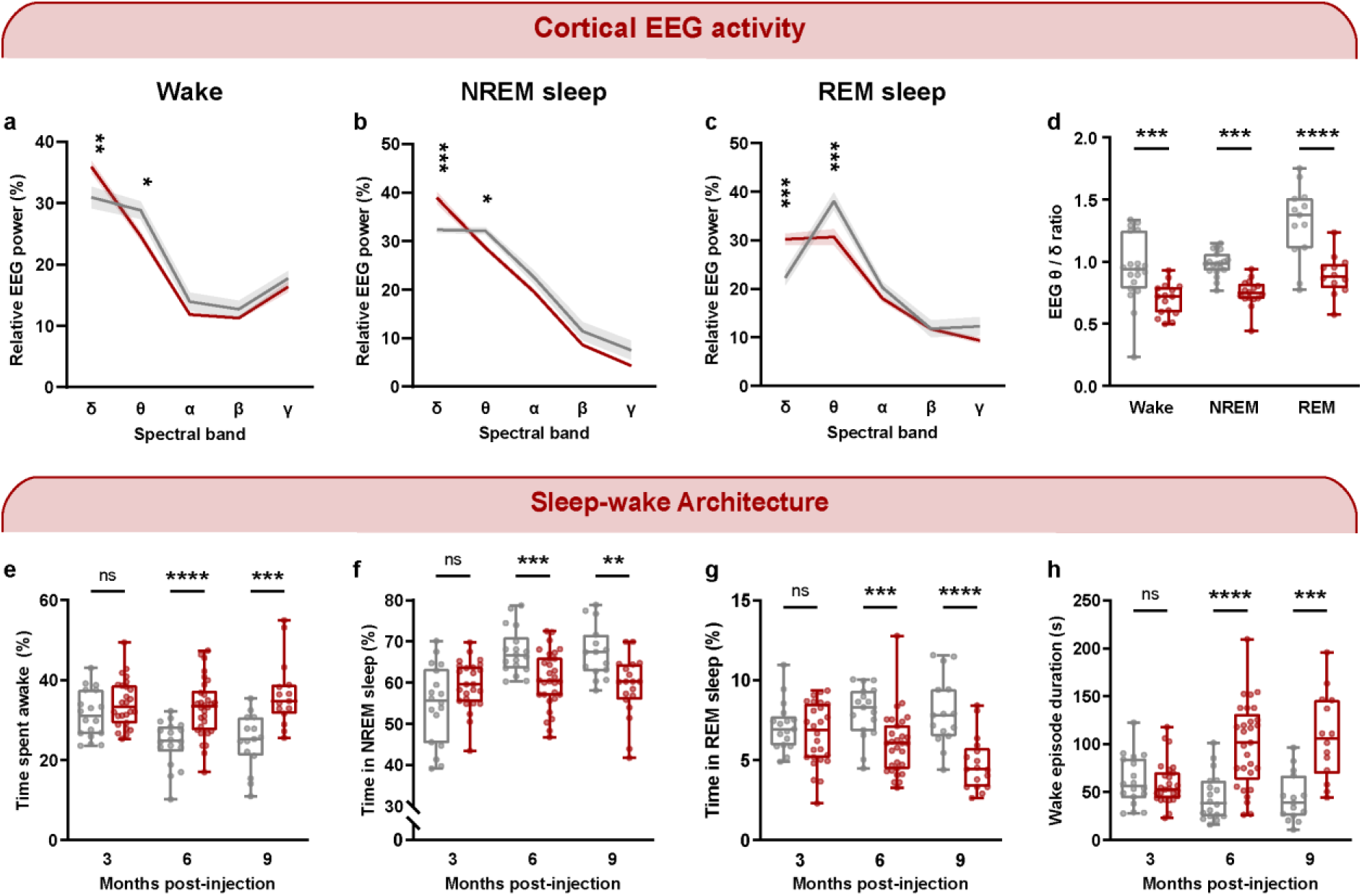
PFF-injected mice exhibited cortical slowing and sleep loss across sleep-wake states following the initial onset of RBD-like behaviours. **(a-c**) EEG power spectrum distribution for PFF-injected mice (red) and PBS controls (grey) at 6mpi during wakefulness (**a**), NREM sleep (**b**), and REM sleep (**c**) (\**P* < 0.05, \*\**P* < 0.01, \*\*\**P* < 0.001, *n* = 16-18). Note that PFF-injection caused a significant reduction in theta power and increased delta power across all states. (**d**) Consequently, the ratio of EEG spectral power between the theta and delta bands was significantly reduced across all sleep-wake states at 6 months post-PFF injection (\*\*\**P* < 0.001, \*\*\*\**P* < 0.0001, *n* = 16-18). (**e-g**) From 6-9mpi, mice spent more time awake during the recording interval (**e**), resulting in corresponding decreases in the overall amounts of NREM (**f**) and REM sleep (**g**) (\*\**P* < 0.01, \*\*\**P* < 0.001, \*\*\*\**P* < 0.0001, *n* = 18-22). (**h**) These effects of PFF injection on sleep-wake architecture were likely driven by prolonged wake episodes at 6 and 9mpi (\*\*\**P* < 0.001, \*\*\*\**P* < 0.0001, *n* = 18-22).

### Development of motor deficits in mice injected with αSyn PFF into the SLD

Wildtype mice that received unilateral PFF or PBS injections also underwent a longitudinal battery of motor tests (at 3, 6, and 12 mpi) to determine whether the propagation of synucleinopathy beyond the SLD led to the development of PD-like motor deficits during wakefulness. The testing battery comprised of open field, rotarod, grip strength, and catwalk gait analysis. The latter quantifies various spatiotemporal parameters of gait (e.g., stride length, swing duration, and paw placement) by capturing and analyzing each animal’s footprints as it voluntarily traverses a glass walkway (Fig. 5a). No motor impairments were detected prior to 6 mpi in PFF– or PBS-injected mice. However, PFF-injected mice developed significant gait alterations relative to age-matched control animals at 12 mpi. Specifically, PFF-injected mice displayed prolonged swing speed and increased time of single stance in both forelimbs and hindlimbs (Fig. 5b). Unlike the loss of REM sleep muscle atonia which occurred in both males and females, gait abnormalities were evident only in male mice injected with PFFs (Supplementary Fig. 6a), with females showing no significant difference from controls (Supplementary Fig. 6b). In contrast, mice inoculated with PFFs did not differ in open field, grip strength, or rotarod performance at all time points examined in this study (Supplementary Fig. 7a-c). Similarly, there were no differences between PFF-injected and control mice across several cognitive tasks assessed at 12 mpi, including Y-maze, novel object recognition, and fear conditioning tests (Supplementary Fig. 7d-f).

**Fig. 5.**
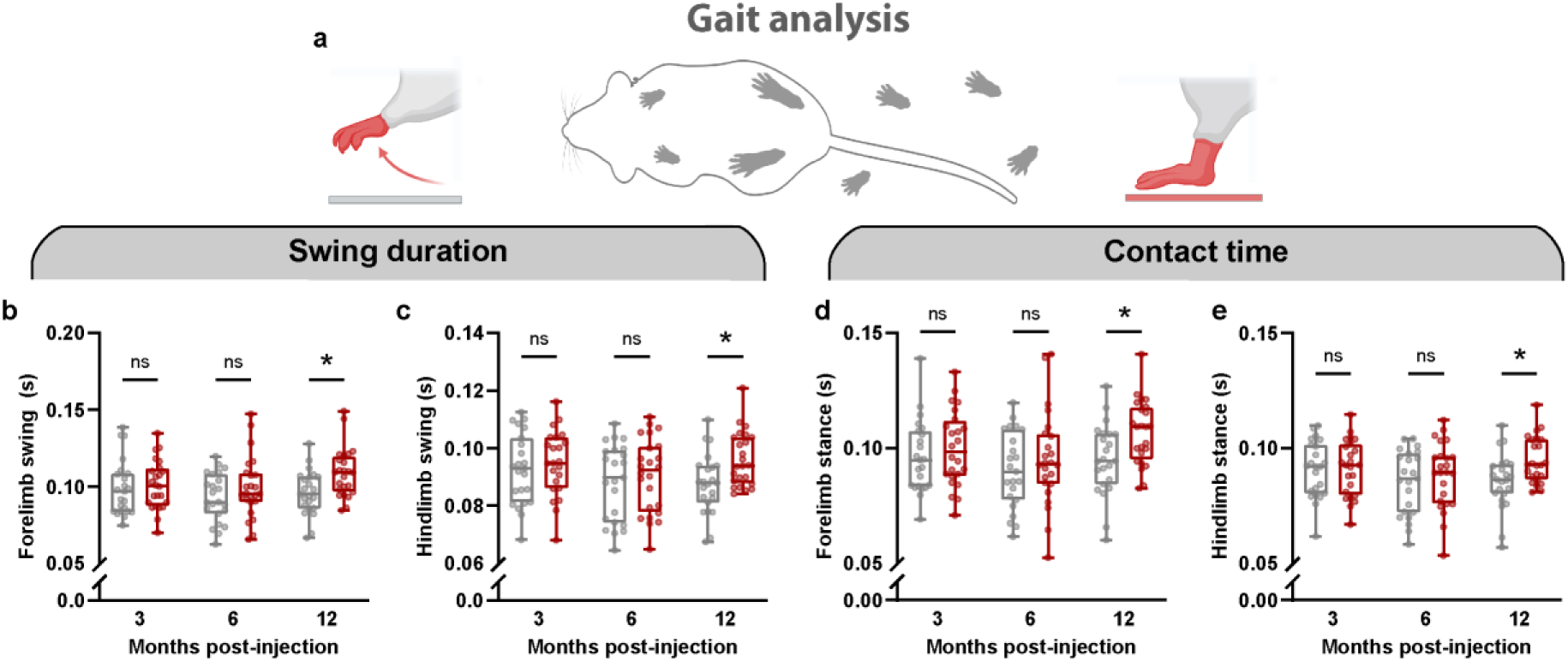
Propagative ɑSyn pathology was associated with gait dysfunction. The CatWalk XT gait analysis system was used to analyze the gait of PFF-injected mice from 3-12mpi. Stance and swing measures were recorded from forelimb and hindlimb pairs, and the average was taken between left and right limbs from each pair. (**a**-**b)** PFF-injected mice developed prolonged swing in both forelimbs (**a**) and hindlimbs (**b**) at 12 mpi (\**P* < 0.05, *n* = 23-24). (**c**-**d**) PFF-injected mice developed contact time in both forelimbs (**c**) and hindlimbs (**d**) at 12 mpi (\**P* < 0.05, *n* = 23-24).

### Selective vulnerability of SLD/vLDT cellular populations

To further understand the relationship between SLD αSyn pathology, RBD symptoms, and gait deficits in our model, we examined whether certain neuronal subpopulations in this region were disproportionally affected by PFF-induced pathological seeding as each behavioural phenotype emerged. Given that PFF-induced αSyn pathology affected both the SLD and the neighboring vLDT, we quantified αSyn aggregates and neuronal loss within a broader region in the dorsal pons that encompassed both nuclei (i.e., the SLD/vLDT complex). Sections at the level of the SLD/vLDT were double-labeled for p-αSyn and one of the major neurotransmitter phenotypes found in the pons (glutamate, GABA, noradrenaline, or acetylcholine). At 1 mpi, the majority (>80%) of αSyn pathology-bearing cells colocalized with choline acetyltransferase (ChAT) while the remaining p-αSyn positive neurons expressed the vesicular glutamate transporter vGlut2 (Fig. 6a-d). Neurons expressing vGAT or tyrosine-hydroxylase represented <1% of all αSyn aggregate-bearing cells.

**Fig. 6.**
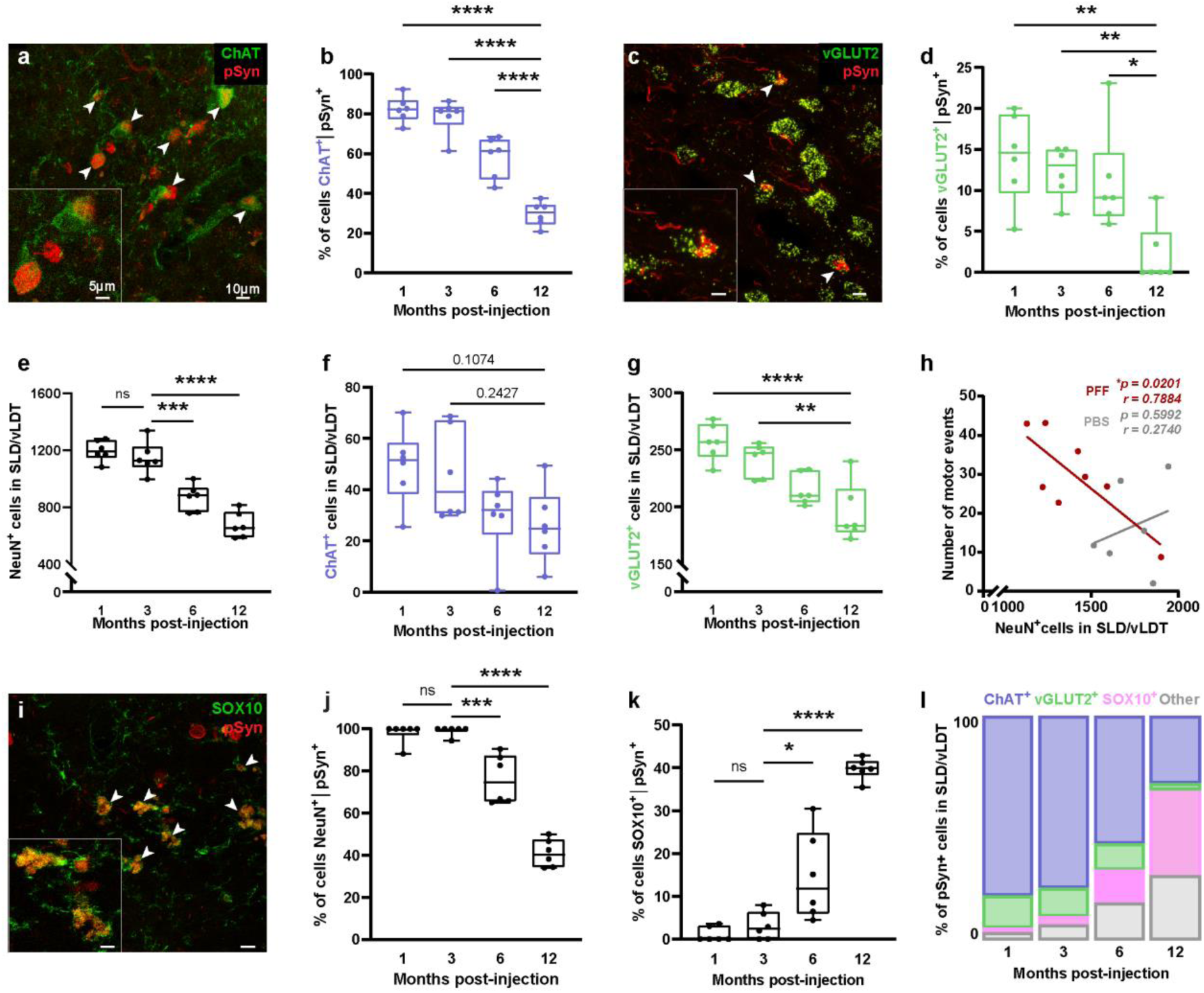
αSyn PFFs injection seeded intracellular aggregation of αSyn in diverse cell types and induced degeneration in the SLD/vLDT. **(a**) Representative image shows immunostaining for p-αSyn (red) and ChAT (green) in the SLD/vLDT from a αSyn PFF-injected mouse at 9 mpi. p-αSyn-positive aggregates were colocalized within ChAT+ neurons (arrowheads). (**b**) Cholinergic neurons in the SLD/vLDT accounted for the majority of aggregate-bearing cells per section until 3 mpi. From 6 mpi, the proportion of p-αSyn-positive aggregates in cholinergic neurons gradually decreased. (**c**) Representative micrograph showing colocalization of p-αSyn immunostaining (red) with vGLUT2 mRNA. (**d**) Glutamatergic SLD/vLDT neurons accounted for a relatively small proportion of all aggregate-bearing cells per section, which decreased at 12 mpi. (**e-g**) Quantification of the total number of NeuN– (**e**), ChAT– (**f**) and vGLUT2-positive cells (**g**) in the SLD/vLDT. A progressive decrease in NeuN positive cells is observed from 1-12 mpi, which was largely driven by a reduction in glutamatergic SLD/vLDT neurons. (**h**) Linear regression analysis reveals a negative correlation between the total number of SLD/vLDT neurons and RBD-like behaviours in PFF-injected mice (red) but not in PBS controls (grey). Mice with the fewest remaining neurons (more severe degeneration) exhibited the highest levels of motor activity during REM sleep. (**i**) Representative micrograph shows immunostaining for p-αSyn (red) and SOX10 (green) in the SLD/vLDT. (**j**-**k)** From 1-12mpi, there was a decrease in the proportion of aggregate-bearing cells within neurons (**j**), and an increase in the proportion of aggregate-bearing oligodendrocytes (**k**) in the SLD/vLDT. (**l**) Diagram shows the proportion of all aggregate-bearing SLD/vLDT cells that co-express ChAT, vGLUT2 or SOX10 in PFF-injected mice from 1-12mpi.

The proportion of p-αSyn positive cells containing ChAT was reduced to ∼60% by 6 mpi and progressively declined to ∼30% at 12 mpi. The proportion of p-αSyn containing cells expressing vGlut2 also decreased over time, with a ∼80% reduction in glutamatergic neuron pathology occurring between 6 to 12 mpi. Notably, the total number of neurons in the SLD/vLDT also decreased over the time points examined, with nearly half of the original number of neurons lost by 12 mpi (Fig. 6e and Supplementary Fig. 1a-c). Quantification of the total number of ChAT and vGlut2 neurons showed that the former remained stable over time (Fig. 6f) whereas a significant reduction in the number of glutamatergic neurons in the SLD/vLDT was observed from 1 to 12 mpi (Fig. 6g).

Linear regression analysis of EMG measurements paired with SLD/vLDT neuronal counts from individual PFF-injected animals revealed that the number of surviving SLD/vLDT neurons was inversely correlated with RBD-like behaviours in PFF-injected mice but not control animals (Fig. 6h), such that mice with the fewest remaining SLD-/vLDT neurons had the highest motor activity in REM sleep. These results support a direct relationship between the burden of αSyn pathology, neurodegeneration and RBD symptom severity, consistent with previous clinical imaging findings^38^ and the physiological role of SLD/vLDT neurons in REM sleep motor suppression. We next assessed the contribution of specific SLD/vLDT neuronal subtypes to RBD by performing a similar analysis for glutamatergic and cholinergic cells, the main neuronal populations affected by PFF-induced synucleinopathy. We observed a strong negative correlation between the number of glutamate cells affected by αSyn pathology and RBD-like behaviours (Supplementary Fig. 8a), whereas no such correlation was observed for cholinergic neurons (Supplementary Fig. 8b). Taken together, the degree of SLD/vLDT glutamatergic neuronal loss was predictive of RBD symptom severity.

Despite significant neuronal loss, large numbers of p-αSyn positive cells continued to persist in the SLD/vLDT up to 12 mpi, prompting us to examine whether other cell types also developed αSyn inclusions in this region. While the number of astrocytes and Iba-1-positive microglia increased in the SLD/vLDT of PFF-injected mice at 9 mpi (Supplementary Fig. 9), they were devoid of p-αSyn immunoreactivity. In contrast, inclusion-bearing oligodendroglia labeled by the transcription factor SOX10 were detected at the time when RBD phenotype emerged (Fig. 6i). When both neuronal and oligodendroglia pathology was quantified over time, we found that αSyn pathology was almost exclusively detected in neurons at 3 mpi while the decline in neuronal pathology at 6mpi coincided with a dramatic increase in p-αSyn positive oligodendrocytes (Fig. 6j) which were rarely observed before this time but accounted for ∼40% of total pathology by 12 mpi (Fig. 6k and 6l).

## DISCUSSION

Idiopathic RBD is an established prodromal manifestation of α-synucleinopathies that precedes overt motor and cognitive symptoms in a considerable proportion of subjects. The mesopontine tegmentum and ventral medulla have been identified as regions that regulate REM sleep muscle atonia and therefore represent the likely anatomical substrates for RBD. While post-mortem studies indicate a close association between the presence of αSyn pathology within these brainstem structures and iRBD, this link remains poorly defined largely owing to a lack of clinically-relevant animal models of RBD. Here, we show that the accumulation of abnormal αSyn in the murine mesopontine tegmentum (i.e., SLD) induced by either viral-mediated overexpression or stereotactic injection of αSyn PFFs both led to a progressive phenotype that recapitulated key features of human iRBD. These findings support the functional role of the SLD in REM sleep atonia and implicates this region in the pathogenesis of RBD and progression to other disease symptoms.

In humans, current diagnostic criteria defines RBD as involving phasic and/or tonic muscle hyperactivity during REM sleep^10^. Although previous rodent models of RBD have primarily described increases in nuchal muscle tone accompanied by motor behaviours^16,17,26^, relatively few studies have characterised changes in phasic motor activity during REM sleep. This is important to consider in modeling RBD because phasic motor events tend to be much more common than tonic events in RBD episodes^39,40^. Our detailed EMG analyses bridge this gap between existing animal models and clinical presentation by demonstrating that mice injected with αSyn PFFs can develop both enhanced variability in muscle tone and prominent phasic motor events during REM sleep. Furthermore, recent clinical observations have shown that phasic, but not tonic, elevations in EMG activity effectively differentiate PD patients with RBD from those without and from healthy controls^41^. Notably, phasic RBD metrics have also been associated with magnetic susceptibility changes in the substantia nigra, a correlation not observed with tonic parameters^42^, suggesting that specific RBD features may reflect distinct neuropathological processes. These insights highlight the translational value of replicating phasic motor abnormalities in preclinical models to better mirror the human condition.

To initiate propagative synucleinopathy in the SLD, we injected wild-type mice with αSyn PFFs and characterised behavioural, electrophysiological and histopathological outcomes in these animals longitudinally. These experiments revealed three important findings concerning RBD mechanisms (Fig. 7): First, the SLD and adjacent vLDT are highly permissive to pathological seeding, with glutamatergic and cholinergic neurons exhibiting a markedly heightened susceptibility to the formation of pathology and subsequent neuronal loss. Unexpectedly, we also observed the delayed appearance of significant oligodendrocytic pathology several months after pathology was first seen in neighboring neurons. Second, αSyn pathology in SLD neurons is sufficient to elicit RBD-like behaviours in mice, with the extent of neuronal loss being tightly correlated with the severity of the RBD phenotype. Third, αSyn pathology originating in the SLD spreads to interconnected circuits distributed across the neuraxis, leading to the progressive disruption of multiple aspects of sleep architecture and specific motor impairments. Altogether, these findings provide novel insights into the pathogenesis of RBD at the circuit and phenotypic level.

**Fig. 7.**
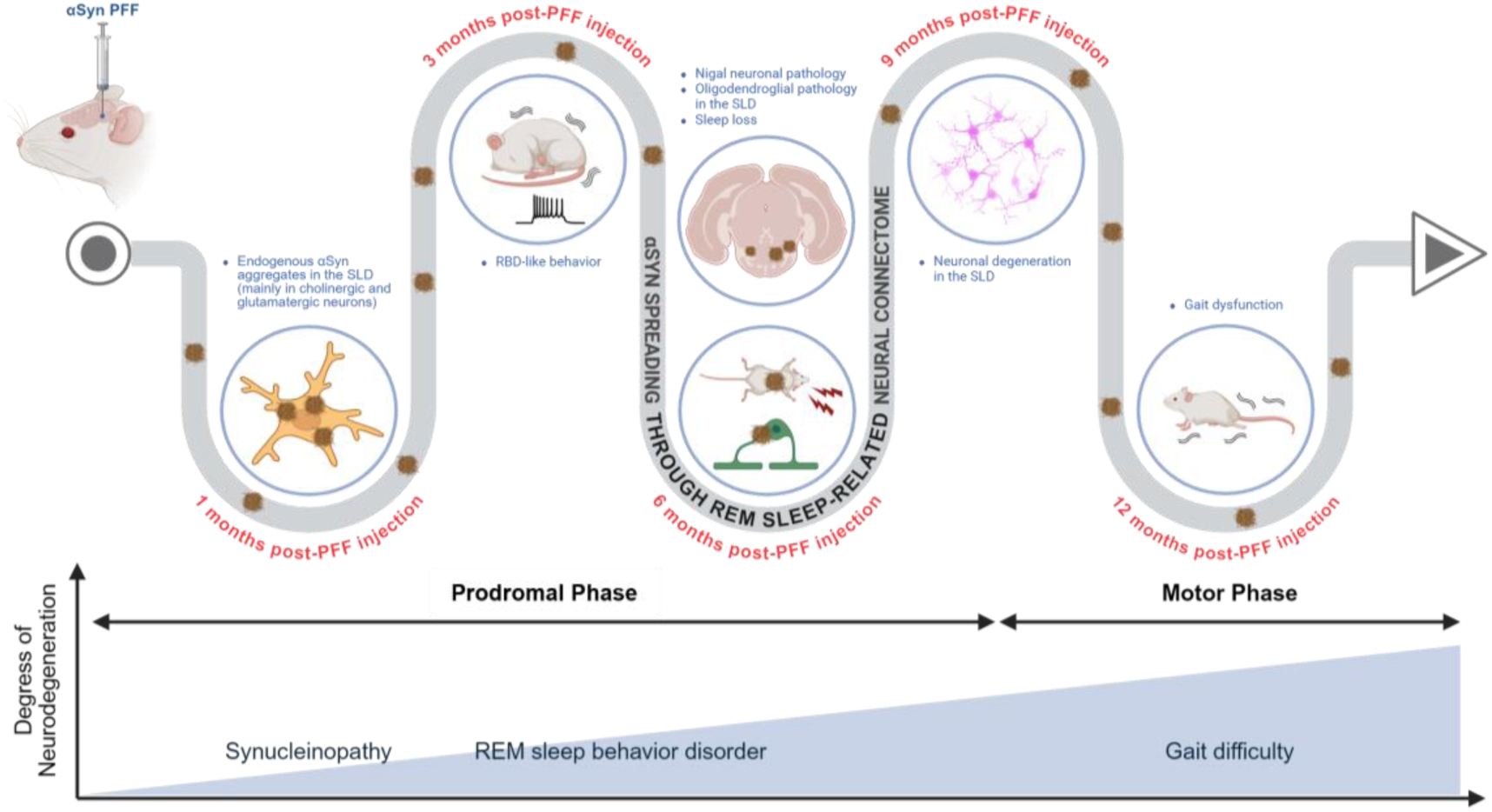
Schematic summary of the temporal progression of αSyn pathology and associated behavioural phenotypes following PFF injection. Following intracerebral injection of αSyn PFFs into mice, endogenous αSyn aggregation (brown, thread-like entangled aggregates) is first observed at 1 mpi in cholinergic and glutamatergic neurons of the SLD/vLDT. Subsequently, αSyn spreads through the REM sleep connectome, coinciding with the emergence of RBD-like behaviours at 3 mpi. By 6 months, αSyn aggregates reach the substantia nigra, and oligodendroglial αSyn pathology accumulates in the SLD. This stage is also accompanied by sleep disturbances. Neuronal degeneration becomes evident at 9 months, and by 12 months, motor impairments including gait dysfunction are observed. The bottom panel illustrates a progressive trajectory from prodromal to motor phases, showing an increasing degree of neurodegeneration and associated symptomatology, including synucleinopathy, RBD-like behaviour, and gait difficulty.

In line with previous post-mortem studies showing that individuals with RBD exhibit αSyn pathology in the SubC^7,20^, we found that this region was affected in a subset of RBD-positive LBD patients. Furthermore, RBD-negative LBD patients displayed only sparse αSyn aggregates in the SubC, suggesting that RBD-positive individuals represent a distinct LBD subtype characterised by mesopontine pathology^43–45^. These clinical findings are consistent with our PFF-based mouse model, where pathology was found in SLD neurons.

SLD glutamatergic neurons are regarded as the principal source of REM sleep atonia generation and maintenance^25^, as the targeted ablation or genetic inactivation of these neurons, or glutamate receptor blockade in this region reliably elicits RBD-like behaviours in animals^26,27,46^. This notion is further supported by recent post-mortem findings from PD patients with RBD revealing the selective loss of SubC neurons that express corticotrophin-releasing hormone binding protein, which are capable of co-releasing glutamate^18^. Neuroimaging evidence from PD patients with RBD also reveal widespread changes in glutamatergic neurotransmission in these individuals compared to RBD-negative individuals^47^. Consistent with these clinical findings, we demonstrate that selective overexpression of ɑSyn in glutamatergic SLD neurons causes RBD-like behaviours in mice and the extent of glutamatergic neuronal loss strongly correlated with the severity of RBD symptoms in PFF-injected animals. Collectively, these results highlight a central role of glutamatergic mespontine pathology in RBD pathophysiology.

Previous clinical data suggest that cholinergic mechanisms may also contribute to RBD. For example, iRBD is associated with cortical cholinergic degeneration of the nucleus basalis of Meynert^48^ and abnormal activity in the pedunculopontine nucleus^49^. Moreover, treatment with acetylcholinesterase inhibitors have been reported to ameliorate RBD symptoms^50^. Our results partially align with these findings, as the predominance of ɑSyn pathology in cholinergic neurons in the SLD/vLDT coincided with the onset of RBD-like behaviours at 3 mpi. However, unlike glutamatergic pathology, no appreciable correlation was observed between cholinergic cell numbers and the severity of RBD-like behaviours in PFF-injected mice. Cholinergic dysfunction may therefore participate in a more modulatory capacity in RBD pathogenesis. Alternatively, cholinergic mechanisms may be more closely linked to other prodromal non-motor features such as cortical slowing or mild cognitive impairment^48^, which frequently co-occur with RBD in prodromal synucleinopathy. Supporting this view, previous animal experiments show that although cholinergic stimulation of the SLD promotes REM sleep, neither selective lesions of pontine cholinergic neurons, nor application of cholinergic receptor antagonists in the SLD substantially alter REM sleep or its component motor atonia^16,51^. Therefore, these observations suggest that SLD glutamate neurons are the primary drivers of RBD symptoms, which could be further modulated by pre– or post-synaptic cholinergic mechanisms. Dysregulation of both glutamatergic and cholinergic signaling in the mesopontime tegmentum may synergistically contribute to RBD pathophysiology, representing a unified and circuit-based framework for RBD that requires further investigation.

As the pathological burden in SLD/vLDT neurons peaked, we observed a dramatic rise in αSyn pathology within oligodendrocytes. Glial cytoplasmic αSyn inclusions are the pathological hallmarks of multiple system atrophy, a synucleinopathy characterised by pronounced pontine atrophy, in which >90% of patients exhibit RBD^52^. Some studies indicate that sleep, especially the REM phase, regulates the proliferation of oligodendrocyte precursor cells and myelination^53^. Moreover, acetylcholine signaling is associated with oligodendrocyte survival, differentiation, and remyelination^54–56^, raising the possibility that early pontine cholinergic deficits increase the vulnerability of oligodendrocytes in the SLD to αSyn pathology. Whether neuron-to-glia transfer αSyn pathology occurs in this context and oligodendroglial dysfunction contributes to RBD symptomology remains to be resolved.

Whole-brain histopathological analysis of our PFF-injected cohorts showed that synucleinopathy expands through the SLD connectome. At the earliest time point examined (1 mpi), pathology in the SLD propagated to structures directly regulating REM sleep in the brainstem including the ventral medulla, pedunculopontine nucleus, parabrachial nucleus, and periaqueductal gray, likely underlying the phasic REM sleep muscle activity that manifested by 3 mpi. By 6-12 mpi, synucleinopathy had infiltrated neural structures related to wakefulness, such as the lateral hypothalamus, tuberomammillary nucleus, dorsal raphe, corresponding to the observed sleep loss and increased wakefulness in these mice.

Besides RBD, sleep disturbances, including insomnia or excessive daytime somnolence, are common and often debilitating non-motor symptoms in patients with PD^34–37^. PFF-injected mice exhibited sleep loss due to prolonged wake bouts, consistent with insomnia in PD (i.e., difficulty entering or maintaining sleep)^37^. Similarly, pathology also spread to the limbic system including the amygdala, bed nucleus of the stria terminalis, nucleus accumbens, infralimbic area, and anterior cingulate area, consistent with the emotional component of the dream-enacting behaviours described in RBD patients. The central nucleus of the amygdala is thought to promote REM sleep by inhibiting vlPAG neurons that suppress SLD neurons^57^, while dopamine release in the basolateral amygdala or nucleus accumbens promotes REM sleep^58^. In addition, a FDG-PET study reported significantly increased glucose utilization in limbic and paralimbic areas during REM sleep relative to waking^59^. Collectively, our findings suggest that αSyn propagation through the neural connectome faithfully reproduces prodromal sleep disturbances.

Gait disturbances were the primary motor manifestation observed in PFF-injected mice, which coincided with the propagation of αSyn pathology from the SLD to more distal components of the SLD connectome. These motor deficits were not the result of reduced muscle strength, as grip strength remained comparable to that of controls. Interestingly, mice injected with PFFs performed similarly to controls in the rotarod test, which requires sustained high-speed locomotion, suggesting that the challenge may lack the sensitivity to detect subtle impairments in gait^60^. A longitudinal study which followed patients with iRBD for up to 12 years documented that gait speed was reduced approximately 6.5 years before phenoconversion^3^. Although mild axial and limb bradykinesia preceded gait impairment before phenoconversion, these symptoms are difficult to detect objectively in rodents. Using a detailed analysis of ∼50 gait-related parameters, we observed significant changes in swing duration and contact time for both fore– and hindlimb during locomotion at 12 mpi, reflecting a prolonged latency in the execution of a forward step (i.e., delayed lifting and advancement of a single limb). In line with our mouse model, patients with RBD also show increased variability in their step width, more asymmetry in step length, and decreased velocity/cadence^61^. Since nigral pathology was not prominent in our mouse model, αSyn pathology in the pedunculopontine nucleus or midbrain reticular nucleus may better account for the observed prodromal gait abnormalities in our model^62,63^.

In summary, our study presents an in vivo model framework that links αSyn pathology in the SLD to the emergence of a progressive RBD-like phenotype and subsequent motor dysfunction. We demonstrate that αSyn aggregates initially accumulate in cholinergic and glutamatergic neurons of the mesopontine tegmentum, coinciding with the onset of RBD-like behaviours. As pathology propagates throughout the CNS, including additional midbrain, hypothalamic and forebrain structures, animals progress from an iRBD phenotype characterised by motor dysfunction during REM sleep to also developing waking motor deficits and broader disruptions in sleep-wake architecture that are characteristic of clinical PD. These findings highlight the SLD as an early and vulnerable hub in the prodromal phase of LBD and other synucleinopathies, providing novel insights into the mechanisms that drive disease progression throughout the course of the synucleinopathies.

## METHODS

### Human patient samples

Post-mortem histopathological assessment was conducted on formalin-fixed, paraffin-embedded brainstem sections from individuals with PD with comorbid RBD, PD without RBD and healthy individuals. All brain samples were recovered from the University of Pennsylvania Center for Neurodegenerative Disease Research (CNDR) Brain Bank^64^. First, we enrolled 20 cases who (1) were clinical diagnosed as Parkinson’s disease, Parkinson’s disease with dementia, or dementia with Lewy bodies, (2) were neuropathological diagnosed as Lewy body disease, and (3) had the information on RBD. The presence of RBD was determined by reporting dream enactment behaviour from the patient’s bed partner and either an RBD screening questionnaire score of 7 or higher^65^ or confirmation by video-polysomnography. After excluding cases who had a borderline RBD screening questionnaire score (4 to 6) or had a poor quality of autopsy sample, we finally enrolled 10 LBD cases and grouped 6 cases as the RBD negative (LBD-RBD-) group and 4 cases as the RBD positive (LBD-RBD+) groups. We also enrolled in 4 control cases whose primary neuropathological diagnosis was either unremarkable or primary age-related tauopathy. Detailed demographic information on these patients was described in Supplementary Table 1. Area of the SubC was confirmed by human brainstem atlas^66^ and neuronal staining using the antibody for the tyrosine hydroxylase.

The use of postmortem tissue was approved by the Institutional Review Boards of the University of Pennsylvania. Patients or their next of kin provided written informed consent for autopsy and the use of tissue samples.

### Animals

All experimental protocols were jointly approved by Institutional Animal Care and Use Committees at the University of Pennsylvania and University of Toronto. Procedures were conducted in accordance with NIH and Canadian guidelines on animal care. Male and female wild-type mice of C57BL/6/ (IMSR_JAX:000664) or B6C3F1/J (IMSR_JAX:100010) genetic background were used for AAV-ɑsyn and PFF injections, respectively. To allow for the selective overexpression of ɑSyn in glutamate SLD/vLDT neurons, we procured transgenic Vglut2-ires-cre knock-in mice (i.e. Slc17a6tm2(cre)Lowl on C57BL/6 background, Jackson Laboratory). Mice were genotyped using PCR with Cre-recombinase primers (Cre forward: CAC GAC CAA GTG ACA GCA AT; Cre reverse: AGA GAC GGA AAT CCA TCG CT). To account for confounding effects of age or biological sex, experiments were performed on young adult mice of both sexes (age: 10 ± 4 weeks; mass: 23 ± 3g depending on sex). Food and drinking water were provided ad libitum and mice were maintained on a 12:12 light-dark cycle (lights on 07:00-19:00).

### Study design for animal experiments

Stereotaxic inoculation of recombinant αSyn PFF or PBS was performed in the CNDR at the University of Pennsylvania. After surgery, mice were grouped into three cohorts for histopathological, electrophysiological, and behavioural studies, respectively (Supplementary Fig. 10). Bilateral PFF inoculations for the electrophysiological studies were carried out at the University of Toronto. Histopathological analyses were performed at 1, 3, 6, and 12 months post-PFF injection in the CNDR at the University of Pennsylvania to assess the spread of αSyn pathology throughout the brain and identify cellular subpopulation that are vulnerable to αSyn pathology within the SLD. Electrophysiological studies were conducted at 3, 6, and 9 months post-PFFs or PBS injection at the University of Toronto to identify the development of RBD phenotypes and quantify RBD-related parameters as well as sleep architecture. After finishing electrophysiological evaluation at 9 mpi, the mice in the electrophysiological cohort were sacrificed to perform histopathological analyses. Behavioural studies were performed at 3, 6, and 12 months post-PFFs or PBS injection in the Neurobehaviour Testing Core at the University of Pennsylvania to investigate changes in motor function.

### Mouse recombinant αSyn purification and *in vitro* PFFs generation

Wild-type recombinant αSyn proteins production, purification, and fibrilization were conducted as previously described^67^. In brief, the plasmid encoding a full-length mouse αSyn gene were introduced into BL21 (DE3) RIL-competent E. coli (Agilent Technologies #230245) and proteins were purified following lysis using size exclusion and anion exchange chromatography. Purified protein (monomer) was concentrated to 5 mg/ml and kept frozen until use. Recombinant αSyn PFFs were produced by shaking 500 μL of recombinant αSyn monomer (5 mg/ml, 360 μM) in a thermomixer (Eppendorf) at 1,000 rpm at 37 C° for 7 days. Fibril formation was verified by electron microscopy, sedimentation assay, and Thioflavin S staining as described^68^. Before conducting stereotaxic injections, PFFs underwent sonication for 10 cycles (30 seconds ON, 30 seconds OFF, 10°C, high intensity setting) using a bath sonicator (Diagenode Bioruptor Plus or QSonica Q125 with cup horn attachment) to ensure fibrils were of appropriate size to elicit a pathogenic response in vivo.

### Anatomical delineation of the SLD/vLDT complex

The SLD is anatomically defined as the region enclosed by the LDT, parabrachial nucleus, and motor trigeminal nucleus. This definition is broadly consistent with previous functional studies that have identified the REM sleep active neurons in this region through immediate early gene expression, unit recording and fiber photometry, as well as retrograde tracing studies from the ventral medial medulla or the spinal cord ventral horn^16–18,26,69,70^. However, slight variation in SLD boundaries exist across previous studies and mouse brain atlases. In-situ hybridization data from the Allen brain atlas indicates *Chat* mRNA is abundantly expressed in the dorsomedial part of the region corresponding to the vLDT in other atlases. Meanwhile, *Vglut2* is highly expressed in the ventrolateral part of the region, corresponding to the dorsal aspect of the subcoeruleus nucleus in the Paxinos & Fraklin atlas. To account for these discrepancies in anatomy, p-αSyn aggregates and neuronal counts were quantified within a broader region in the mesopontine tegmentum that included both the SLD and the vLDT.

### AAV materials

Two AAVs were used to drive the overexpression of αSyn in SLD neurons. First, an AAV2 serotype was employed to induce the overexpression of wild-type human αSyn in all SLD neurons, using the chicken beta-actin promoter (AAV-ɑSyn; construct: AAV2-CBA-ɑSyn). To restrict ɑSyn overexpression to glutamate SLD neurons, we used a previously characterised AAV9 serotype packaged with the sequence encoding human A53T ɑSyn flanked by a pair of loxP sites^71^. Cre-inducible transduction of ɑSyn in glutamate cells was achieved upon virus injection into Vglut2-ires-cre transgenic mice, which selectively express cre-recombinase in Vglut2-expressing neurons. Viral transgene expression was driven under the elongated factor 1ɑ promoter and enhanced by the woodchuck post-regulatory enhancer (AAV-DIO-ɑSyn; construct: AAV9-Ef1a-DIO-A53T-ɑSyn-WPRE). Control mice were injected with viruses expressing inert fluorescent proteins, either GFP or eYFP (AAV-GFP or AAV-DIO-eYFP). However, due to high attrition in the latter group and limited sample size, we compared both cell-specific and nonspecific αSyn overexpression experiments with the AAV-GFP controls. AAV-ɑSyn was acquired from the Michael J. Fox Foundation and AAV-DIO-ɑSyn was generously provided by Dr. Ronald L. Klein.

### Stereotaxic inoculation of recombinant αSyn PFF and AAVs

Intracerebral injections of AAVs or ɑSyn PFFs into the murine SLD were performed as previously described with minor modifications^27,67^. Adult mice of 2 – 3 months of age were deeply anesthetized and then secured into a stereotaxic frame (David Kopf Instruments, USA), before a craniotomy was performed to expose a 1mm opening in the skull above the SLD. Infusions were carried out using a digital microinjection syringe pump (Harvard apparatus) at a rate of 0.1 μL/min. AAV infusions were performed to a total volume of 0.4 μL (0.2 μL per hemisphere) bilaterally at 0.05 μL/min into the SLD (stereotaxic coordinates: –5.20 mm relative to bregma; +0.9 mm from midline; –4.25 mm from bregma). Separate groups of mice received bilateral infusions of αSyn PFFs into the SLD (0.2 μg / 1μL PFFs per hemisphere), or unilateral infusions into the right SLD (0.5 μg / 2.5μL PFFs). Mice were administered postoperative analgesic and recovered for at least 14 days after the surgery.

### Tissue preparation

Formalin-fixed paraffin-embedded (FFPE) samples were used for the histopathological cohort. At 1, 3, 6, and 12 mpi, mouse brains were perfused intracardially with 15 mL of PBS and 15 ml of 4% (w/v) paraformaldehyde (PFA) in PBS and fixed in 4% (w/v) PFA in PBS at 4°C overnight. The brains were embedded in paraffin and then sectioned with a thickness of 6 μm using a microtome and mounted on glass slides.

Frozen tissue samples were used to conduct histopathological study for the electrophysiological cohort after completion of electrophysiological experiments at 9 mpi. Mice were deeply anesthetized, transcardially perfused with PBS followed by 4% PFA and their brains harvested. Following fixation, brains were cryoprotected with 30% sucrose, frozen in Tissue-Tek O.C.T. compound (Sakura) and cut into 30 µm-thick coronal sections.

### Immunohistochemistry and fluorescence in-situ hybridization of injected mouse brains

Immunohistochemical staining of FFPE samples was performed to see the spreading of p-αSyn pathology. Every 20^th^ paraffin section throughout the brains was incubated at 4°C for 2 days with the following primary antibodies: an anti-phospho-serine129 αSyn (p-αSyn) antibody and an anti-TH antibody. For generating diaminobenzidine (DAB) reaction products, biotinylated secondary antibodies (Vector laboratories #BA-1000, 1:1000) were used and nuclei were counterstained with hematoxylin.

Immunofluorescence staining of FFPE samples was performed to investigate the cellular subpopulation in the SLD/vLDT which was vulnerable to αSyn pathology. The sections were incubated at 4 °C for 2 days with the following primary antibodies: an anti-p-αSyn antibody, an anti-ChAT antibody, an anti-NeuN antibody, an anti-GFAP antibody, an anti-Iba1 antibody, and an anti-SOX10 antibody. Fluorescent dye-conjugated secondary antibodies were used, and nuclei were stained with DAPI.

Immunofluorescence staining of fresh frozen tissue samples was performed on free-floating brain sections as previously described^27^. Sections were washed, blocked in 4% goat serum for 1 hour, then incubated in primary antibodies for up to 48 hours at 4°C. For visualization, sections were incubated in fluorophore-conjugated secondary antibodies for 1 hour at room temperature, counterstained with DAPI, and mounted on glass slides.

RNA-based in-situ hybridization was used to characterise the molecular identity of pathologically affected SLD/vLDT neurons. Mouse brains were collected as described above, cut into 16 µm-thick sections for fresh frozen samples and into 6 µm-thick sections for FFPE samples, then mounted on glass slides. The RNAscope multiplex fluorescent reagent version 2 assay (RNAscope, Advanced Cell Diagnostics) was conducted on brain sections according to manufacturer protocol. Target retrieval reagent and protease were sequentially added to slides, followed by application of the target probe for Slc17a6 (VGLUT2) and Slc32a1 (VGAT) mRNA transcripts. The signal was then amplified with multiplex fluorescent v2 AMP1 solution then visualized with Cy3 fluorescent dye. To stain for αSyn aggregates, p-αSyn immunostaining was performed after the RNAscope protocol on slide-mounted sections.

For immunofluorescence and in-situ hybridization staining of FFPE samples, sections were examined with an Eclipse Ni microscope (Nikon) or using a Lamina Scanner (PerkinElmer; 20 × objective) in the University of Pennsylvania. The number of cellular αSyn pathology was counted manually. The number of neurons or glia was counted using QuPath software (version 0.5.0)^72^. We annotated the area of SLD/vLDT using a brush annotation and detected neurons or glia using ‘cell detection’ function with the following parameters: for setup parameters, detection channel = green, required pixel size = 0.5 m; for nucleus parameters, background radius = 8 μm, median filter radius = 0 μm, sigma = 1.5 μm, minimum area = 10 μm^2^, maximum area = 400 μm^2^; for intensity parameters, threshold = 20, cell expansion = 5 μm.

For fresh frozen tissue samples, both upright (Zeiss AxioImager.Z1) or confocal (Zeiss LSM 900 or Leica TCS SP8) microscopes were used to acquire images of immunofluorescent signals in the University of Toronto. All quantitative image analyses were performed at three rostral-caudal positions of the SLD/vLDT. The same approach was used to quantify immunofluorescent signals in experimental and control animals. To assess neuroinflammatory changes in the SLD/vLDT, we stained for astrocytic and microglial markers, then used an image analysis module in ZEN software (Zeiss) to calculate the percentage of a 1.22 mm x 1.22 mm area containing the SLD/vLDT that was immunoreactive for each glial marker. To assess neuronal loss in the SLD/vLDT, we stained for the neuronal marker NeuN and used an image analysis module (ZEN, Zeiss) to objectively count the number of NeuN-positive cell numbers within a 750 µm x 650 µm area corresponding to the SLD/vLDT. During image analysis, automated errors were corrected by an experienced observer to ensure that image artifacts were not included. Primary and secondary antibodies used in this study are listed in Supplementary Table 2.

### Automatic quantification of αSyn pathology and generation of heatmap

The whole process of automatic quantification for DAB immunohistochemical staining of FFPE samples is provided in Supplementary Fig. S11. Scanned whole-slide images were imported into QuPath to generate two subsets of images in PNG format compatible with a workflow modified from QUINT^73^: “registration” images (low-resolution full-color PNGs for Allen Brain Atlas registration) and “segmentation” images (full-resolution binary PNGs containing only pathology-positive pixels). Registration images were given to DeepSlice^74^, then manually adjusted for increased accuracy. This output was run through the “quantifier” function in NUTIL to generate a regional breakdown of each image^75^. Images where 30% or more of the total area contained ROIs were considered relevant for further processing and the corresponding segmentation images were manually filtered for artifacts in GIMP, which primarily consisted of non-specific edge staining. Filtered segmentation images were then fed back into the NUTIL quantifier function to generate a pathology pixel count for each brain region. An R script was created to compile the data for each brain across the study, calculate load metrics (pathologic area/total region area,) and to filter out irrelevant regions. Where noted, sub-regions defined by the Allen Brain Atlas were combined into their parent features. Resultant data was used to generate heatmaps in GraphPad Prism (version 10).

### Electroencephalographic and electromyographic electrode implantation

For the evaluation of sleep-wake behaviours, mice were instrumented with electroencephalographic (EEG) and electromyographic (EMG) electrodes, which respectively allow for cortical and muscle activity measurements^27^. Electrodes were fashioned with stainless-steel wire, soldered onto a microstrip conductor, and assembled into a headplug. Depending on the experiment, mice were instrumented at 2 weeks post-AAV injection or 10 weeks post-PFF injection, to allow for recording of sleep-wake behaviours at predetermined timepoints. During EEG/EMG instrumentation, animals were prepared as described above and elsewhere^27^, then transferred onto a stereotaxic apparatus under anesthesia. Four craniotomies were performed to allow for attachment of EEG electrodes onto the skull with stainless-steel miniature screws (J.I. Morris), and the headplug implant was then secured using dental cement (C&B Metabond and 3M Ketac Cem). EMG electrodes were sutured into the neck and right masseter muscles. Mice were administered postoperative analgesic and recovered for a minimum of 14 days before behavioural experiments.

### Electrophysiological data acquisition

EEG, EMG, and video data were collected to monitor sleep-wake behaviour and allow for the identification of potential RBD-like behaviours in mice. Following recovery from EEG/EMG implantation surgery, mice were transferred to sound-attenuated recording enclosures and their implants were connected with a lightweight cable to a Physiodata Amplifier system (Grass 15LT, Astro Med.) for data acquisition. Electrophysiological signals were amplified by 500-1000x, digitized at 1000Hz (Spike2 Software, 1401 interface, Cambridge Electronic Design Ltd.) and band-pass filtered (EEG, 0.1-100Hz; EMG, 30-3000 Hz). Prior to data analysis, EMG signals were waveform-rectified and notch-filtered to remove 60Hz electrical noise. Videos were synchronized with EEG/EMG data in Spike2 software.

Virally injected mice were recorded once at 8 – 10 weeks post-AAV injection, whereas PFF-injected mice were recorded at 3, 6 and 9 mpi to allow for longitudinal assessment of RBD-like behaviours over time. Mice were allowed to acclimatize in the recording enclosure for 3 days before sleep-wake behaviours were recorded over 24 hours for 2 consecutive days. On completion of the last day of recordings, mice were disconnected from recording cables then sacrificed for tissue collection or singly housed until their next set of recordings.

### Analysis of sleep-wake behaviours

In RBD patients, symptoms are most prominent in the early morning hours, when REM sleep quantities are greatest. In mice, REM sleep amounts are greatest between 14:00 – 17:00 (ZT 7-10). As such, we focused on our analysis of sleep-wake behaviours during this period. Vigilance states were manually classified into 5-second epochs based on standard criteria^27^ using a custom script in Spike2 (CED). Bouts of REM sleep were identified based on low-amplitude, high-frequency EEG activity peaking in the theta frequency range, accompanied by minimal EMG tone with phasic muscle twitches. An Excel macro was used to quantify the number, length, and proportion of the recording interval spent in each vigilance state. To analyze cortical activity, we applied Fast-Fourier transform to EEG signals and performed power spectrum analysis to determine the relative EEG power expressed within each frequency band.

To assess whether mice developed RBD-like behaviours, we analyzed EMG activity during REM sleep. In human patients, RBD symptoms consist of increases in tonic, phasic, or both components of REM sleep muscle activity^10^, with phasic motor events constituting the vast majority of movements in a typical RBD episode^39,76^. In mice, spontaneous REM sleep is similarly composed of tonic and phasic motor components^27,77,78^. Therefore, to assess RBD-like behaviours in mice, we developed a program based on our previously published methods for quantifying phasic and tonic muscle activity in REM sleep^27,78^. To do this, we used the first 2.25s of REM sleep to set a 99.99th percentile of EMG activity for each episode. Phasic motor events (muscle twitches) were defined as brief EMG activations longer than 5ms that exceeded the 99.99%ile amplitude threshold; tonic activity was equal to or below this threshold. The number and total duration of phasic motor events were quantified for every REM episode from each mouse. To account for variation in EMG electrode placement between mice, REM sleep muscle tone was normalized to the amplitude of the EMG during NREM episodes from the same animal. To examine the temporal profile of movements within individual REM episodes, we standardized episodes by length into 20 subdivisions and quantified movements that occurred in each subdivision along the REM episode.

### Spectral analysis of EEG activity

Power spectrum analysis was performed to evaluate the distribution of EEG power across defined frequency bands. For each mouse, after vigilance states were classified in 5 second epochs, a Fast-Fourier transform (Hamming window, FFT size: 1024 Hz) was applied to all epochs labelled as Wake, NREM sleep or REM sleep for the entire recording period. To normalize EEG signals between mice, the raw power at each frequency is divided by the total EEG power across all frequencies analyzed (0-130Hz), resulting in EEG power for each frequency expressed as a relative proportion of the total spectrum for that mouse. The relative EEG power at each band was then calculated by summing the values for all frequencies within the following ranges: (δ) 0.5-4 Hz, theta (θ) 4-8 Hz, alpha (α) 8-15 Hz, beta (β) 15-32 Hz and gamma (γ) 32-130 Hz. The θ/ δ ratio for each mouse was calculated by dividing by the raw θ power by the raw δ power.

### Behavioural procedures

A behavioural test battery for motor symptoms consisted of the open field activity, accelerating Rotarod, grip strength, and gait analysis and was tested at 3, 6 and 12 mpi in 12 PBS– and 12 αSyn PFF-injected mice. A battery for cognitive symptoms consisted of the Y-maze, novel object recognition, and contextual fear conditioning and was tested at 12 mpi in 6 PBS– and 6 αSyn PFF-injected mice. Prior to these procedures, mice are handled gently for two minutes a day for three days to habituate the mice to the investigator.

### Open field activity

Spontaneous locomotion and rearing activity were assessed in an open field arena. The Photobeam Activity System (San Diego Instruments) was used to acquire data. The clear Plexiglas arena (14 in. x 14 in. x 18 in.) is fitted with a scaffold of IR emitters and detectors to collect peripheral, center and vertical (rearing) beam breaks. After a 30-minute habituation to the testing room, a 10-minute trial began with a mouse placed in the center of the arena. All trials were recorded by high-definition camcorders. Digitally recorded trials were processed for automated analysis by ANYmaze software (Stoelting Co, Il) to obtain additional measures.

### Accelerating Rotarod

The Rotarod (IITC San Diego Ca.) is a one-inch diameter, horizontal rod with a coarse surface, programmable to accelerate at different rates. As the rod accelerates, the mouse must adjust the cadence of its stride to remain on the rod. Three trails per day are performed over three consecutive days with the Rotarod accelerating from four to forty rpm in three hundred seconds. There is a thirty-minute intertrial interval when the mouse rests in its home cage. On the very first trial, the mouse is habituated to the stationary rod for two minutes before rotation begins. A trial ends when the mouse fails to walk, by either falling from the rod or making a full rotation while gripping the rod. Mice that complete the full three hundred-second trial are removed from the rod to end the trial.

### Grip strength

Grip strength meters are used routinely to assess strength as part of a general health assessment. After thirty minutes of habituation to the procedures room, a Grip Strength Meter (IITC Instruments, San Diego, Ca) is used to measure forepaw and hindpaw grip strength on subsequent days. For forepaw, a mouse is lowered by the tail so that it grasps a metal T bar with forepaws only. The mouse is pulled backward slowly by the tail in the horizontal plane to exert a force on the T bar, which is transduced to a meter that records the maximum force (grams) exerted before the release of the bar. Five trials are performed with a twenty-minute intertrial interval. Twenty-four hours after the forepaw trails, hindpaw grip strength is measured. For hindpaw the T bar is replaced with a chicken wire grid. A mouse is lowered so that it grasps the grid with all four paws. Similar to the forepaw test, the mouse is slowly pulled backward in the horizontal plane to exert a force on the T bar which is transduced to a meter that records the maximum force (grams) exerted before the release of the bar. Five trials are performed with a twenty-minute intertrial interval.

### Gait analysis

The Catwalk XT (Noldus Information Technology, Netherlands) is a sophisticated system used to acquire quantitative measures of stride dynamics and paw placement. After a thirty-minute habituation to the darkened procedure room a trial is started by placing a mouse at one end of a forty cm enclosed alley. The mouse walks freely toward a goal box containing its home cage. Data is collected over a twenty cm sampling region. Footfalls are illuminated from below and pixelated to collected data on limb motion, stride dynamics and ambulation patterns. Additionally, the pixelated paw print data provides measures of load, braking, standing and propulsion during passive ambulation. The criteria for a compliant run were set to a duration of one half to ten seconds. At least five runs were collected for each mouse then post hoc visual validation of runs provided at least three runs for each mouse for analysis.

### Y-maze test

The Y-maze is used to assess working memory. Mice were habituated to the procedure room for thirty minutes prior to the test. To start the trial, a mouse was placed at the end of one arm of the maze and allowed to explore for eight minutes. During exploration, mice tend to spontaneously alternate the order of the maze arms that they enter. A mouse was considered to enter an arm when all four paws were in the arm. High-definition recordings of the trials were collected for off-line grading. The percentage of spontaneous alternations (% SA) was calculated using the following formula: % SA = [number of alternations / (total arm entries – 2)] x 100. Reduced spontaneous alternations suggest a lack of familiarity with the arm just exited and indicates a working memory impairment.

### Novel object recognition

The novel object recognition procedure consists of habituation/pre-exposure, acquisition and recall trials. The habituation/pre-exposure trials consist of five-minute trials per day over five consecutive days where the mouse freely explores the empty NOR arena (approx. 1 square foot). Twenty-four hours after the last pre-exposure trial, an acquisition trial was performed. During the acquisition trial, mice were returned to the arena, now containing a pair of the same objects. The object pairs used were either glass bottles or metal bars (2” x 2” x 6”) placed about three inches from the arena walls. Mice freely explore the object pairs for fifteen minutes. The arena and objects were cleaned with 70% EtOH between all trials. Twenty-four hours after acquisition, mice were returned to the arena, with one of the now-familiar objects replaced with a novel object (for example, a bottle may be swapped with a metal bar or vice versa). During the recall trial, mice explore the two objects for 15 minutes. All trials are digitally recorded for automated off-line grading by ANYmaze software (Stoelting Co, Il) Time spent exploring (approaches and sniffing) and visits to each object are determined as the primary dependent variables. Mice show an innate propensity to explore the novel object. Reduced exploration of the novel object suggests reduced familiarity of the objects used in acquisition.

### Context and cued fear conditioning

Mice were handled for an additional day to habituate to the conditioning room. During an acquisition trial, mice were placed in the conditioning chambers within a sound attenuating cabinet (Med Associate, Fairfax, VT). The trial consisted of a 150-second prestimulus epoch before an unsignaled 1.25 mA foot shock was delivered. After a thirty-second post-stimulus epoch, mice were returned to their home cages and colony room. Twenty-four hours after acquisition, mice were placed in the conditioning chamber for a five-min recall trial. Freezing behaviour (no motion except for respiratory movements) was automatically determined by FreezeScan software. An increase in freezing during recall demonstrates that an association of the chamber and the foot shock was made.

### Statistical analysis

Results for all statistical tests are described in the figure captions. Statistical analysis of data and graphing was performed using SPSS 25 (IBM Corp, Armonk, NY), R software (version 4.2.1; R Foundation for Statistical Computing, Vienna, Austria), and GraphPad Prism (version 10, GraphPad Software). For all tests, a critical value of α ≤ 0.05 was set. Outliers were assessed using the ROUT method^79^ with a false discovery rate of 1% used as the exclusion criteria. To compare longitudinal data between two groups, we used a two-way repeated measures ANOVA with Bonferroni post-hoc comparisons, where applicable. In many cases, mice removed their EMG electrodes over time, which led to attrition at later time points. To accommodate for this, we opted to use a non-parametric mixed-effects model for datasets with missing values. To assess changes in muscle activity of REM episodes at the population level, the Kolmogorov-Smirnov test was used to compare the frequency distribution of episodes between groups of mice.

### Data availability

The de-identified data that support the findings of this study are available from the authors upon reasonable request.

## Acknowledgments

We would like to thank the Neurobehaviour Testing Core at UPenn/ITMAT and IDDRC at CHOP/Penn U54 HD086984 for assistance with the behaviour procedures, and Diane Chernoff, Norihito Uemura, Juliana Benitez, Michael Kozak, and Kevt-Her Hoxha for technical assistance.

## Funding

This research was supported by the Michael J. Fox Foundation for Parkinsons’s Research (MJFF-021825 to K.C.L, J.H.P., H.S.Y.), National Institutes for Health (AG084497), Canadian Institutes for Health Research (PJT-159606 to J.H.P), Korean Neurological Association (KNA-20-SK-13 to H.S.Y.), and the Korea Health Technology R&D Project through the Korea Health Industry Development Institute (KHIDI), funded by the Ministry of Health & Welfare, Republic of Korea (grant number: HI19C1330 to H.S.Y.),

## Author contributions

R.L., H.S.Y., J.H.P., J.J.F. and K.C.L. conceived and designed the experiments R.L., H.S.Y., D.M., A.B., C.D., B.J.D., Y.L., B.A., J.A.S., B.Z., A.Y., J.L., and J.J.F performed experiments, and analyzed results. R.L. wrote the paper with input from coauthors. H.S.Y., J.H.P., J.J.F., K.C.L. revised and edited the paper. All authors reviewed and approved the manuscript.

## Competing interests

The authors report no competing interests.

## Supplementary Materials

## Notes

### Competing Interest Statement

The authors have declared no competing interest.

