## Supplementary Tables 1 & 2, Supplementary Figures 1-11 for "Cell vulnerability within the sublaterodorsal tegmental nucleus underlies REM sleep behaviour disorder in prodromal α-synucleinopathy"

### Supplementary Materials

#### Supplementary Table

**Supplementary Table 1.** Demographic characteristics of post-mortem human autopsy samples

**Supplementary Table 2.** List of primary and secondary antibodies used in this study.

#### Supplementary Figures

**Supplementary Fig. 1.** AAV-mediated αSyn overexpression in SLD neurons does not alter REM sleep amounts or EEG activity during REM sleep.

**Supplementary Fig. 2.** PBS injection into the SLD did not cause changes in EMG activity over time

**Supplementary Fig. 3.** Unilateral PFF injection into the SLD induced a moderate and transient increase in EMG activity during REM sleep

**Supplementary Fig. 4.** PFF injection caused no changes in muscle activity during wakefulness or NREM sleep

**Supplementary Fig. 5.** Male and female PFF-injected mice did not exhibit clear sex differences in the expression of RBD-like behaviors

**Supplementary Fig. 6.** Sex differences in gait dysfunction

**Supplementary Fig. 7.** PFF-injected mice did not display severe or cognitive motor deficits

**Supplementary Fig. 8.** Association between neuronal pathological burden and the degree of RBD-like behavior

**Supplementary Fig. 9.** PFF injection induced reactive gliosis in the SLD/vLDT

**Supplementary Fig. 10.** Experiment schematic

**Supplementary Fig. 11.** Pipeline for automatic quantification of p-αSyn pathology and generation of heatmap

**Supplementary Table 1. Demographic characteristics of post-mortem human autopsy samples**

| | Healthy controls | LBD patients without RBD | LBD patients with RBD | *P*-value |
|---|---|---|---|---|
| Number | 4 | 4 | 6 | |
| Age at death, yr | 74.1 ± 7.5 | 81.8 ± 9.7 | 73.1 ± 5.9 | 0.220 |
| Sex, male, *n* (%) | 3 (75.0) | 2 (50.0) | 5 (83.3) | 0.511 |
| Post-mortem interval, month | 19.0 ± 13.5 | 12.8 ± 8.6 | 22.7 ± 18.3 | 0.603 |
| Brain weight, g | 1292.5 ± 318.5 | 1210.0 ± 89.1 | 1381.8 ± 61.8 | 0.354 |

The values are expressed as mean ± standard deviation or number (percentage). *P*-values are results of the independent *t* test for continuous variable and the Fisher's exact test for categorical variable.

**Supplementary Table 1. List of primary and secondary antibodies used in this study.**

| Antigen | Host | Manufacturer | Antigen retrieval | Catalog # | Dilution |
|---|---|---|---|---|---|
| ***University of Pennsylvania*** | | | | | |
| Phosphorylated αSyn (EP1536Y) | Rabbit | Abcam | Sodium citrate | ab51253 | 1:10,000 |
| ChAT | Goat | Sigma Aldrich | Sodium citrate | AB144P | 1:200 |
| NeuN | Mouse | Millipore Sigma | Sodium citrate | MAB377 | 1:200 |
| SOX10 | Goat | R&D Systems | Sodium citrate | AF2864 | 1:3,000 |
| Anti-goat AlexaFluor 488 | Donkey | Thermo-Fisher | NA | A-11055 | 1:500 |
| Anti-mouse AlexaFluor 488 | Donkey | Thermo-Fisher | NA | A-31571 | 1:500 |
| Anti-rabbit AlexaFluor 594 | Donkey | Thermo-Fisher | NA | A-21207 | 1:500 |
| Anti-goat AlexaFluor 647 | Donkey | Thermo-Fisher | NA | A31573 | 1:500 |
| ***University of Toronto*** | | | | | |
| αSyn (LB509) | Mouse | Abcam | - | ab27766 | 1:800 |
| Phosphorylated αSyn (EP1536Y) | Rabbit | Abcam | Sodium citrate | ab51253 | 1:1,600 |
| Phosphorylated αSyn (S129) | Rabbit | Abcam | NA | ab59264 | 1:400 |
| NeuN | Rabbit | Abcam | NA | ab104225 | 1:1000 |
| ChAT | Mouse | Sigma-Aldrich | NA | AMAB91130 | 1:200 |
| SOX10 (20B7) | Mouse | Invitrogen | Sodium citrate | 14-5923-82 | 1:400 |
| Iba1 | Rabbit | Abcam | NA | ab178846 | 1:800 |
| GFAP | Chicken | Abcam | NA | ab4674 | 1:800 |
| Anti-mouse AlexaFluor 488 | Goat | Jackson ImmunoResearch | NA | 115-547-003 | 1:800 |
| Anti-rabbit Cy3 | Goat | Jackson ImmunoResearch | NA | 111-167-003 | 1:800 |
| Anti-chicken AlexaFluor 647 | Donkey | Jackson ImmunoResearch | NA | 703-606-155 | 1:800 |

N/A, not applicable

**Supplementary Fig. 1. AAV-mediated αSyn overexpression in SLD neurons does not alter REM sleep amounts or EEG activity during REM sleep.**

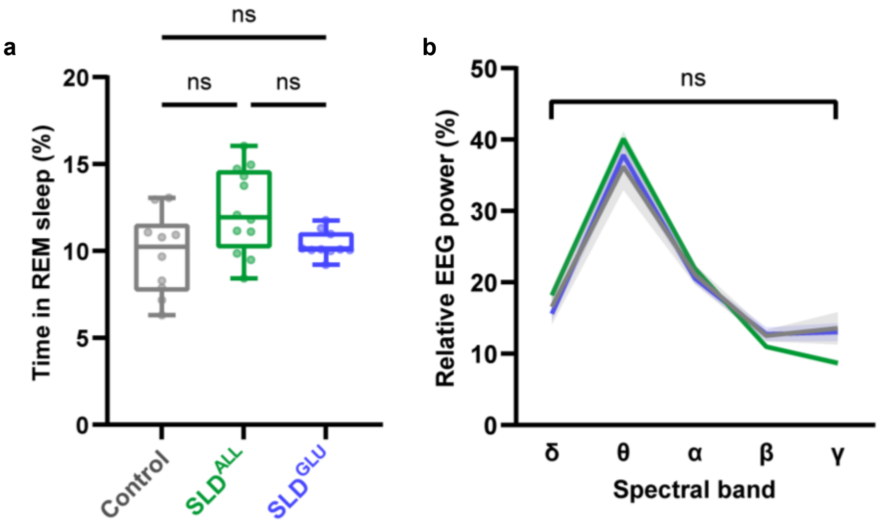


**(a)** At 8-10 weeks post-AAV injection, neither pan-neuronal overexpression of αSyn in the SLD nor cell-specific overexpression in glutamatergic SLD neurons altered the overall amount of REM sleep compared to GFP-transduced control mice ($P > 0.05$, $n = 10\text{-}12$). **(b)** EEG spectral analysis revealed that neither pan-neuronal nor cell-specific overexpression of αSyn in SLD neurons affected cortical EEG activity during REM sleep ($P > 0.05$, $n = 10\text{-}12$).

**Supplementary Fig. 2. PBS injection into the SLD did not cause changes in EMG activity over time.**

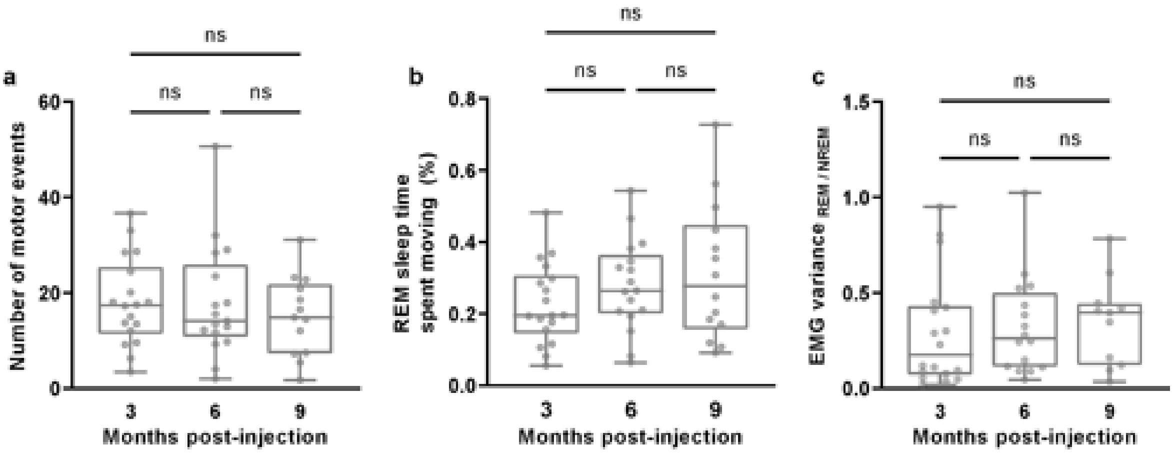


**(a-c)** Throughout the duration of the study from 3-9mpi, PBS injected mice do not show differences in the number of motor events during REM sleep (**a**), the proportion of time that such movements occupy during REM sleep (**b**), nor the REM:NREM ratio of EMG variance (**c**) ($P > 0.05$, $n = 18$).

**Supplementary Fig. 3. Unilateral PFF injection into the SLD induced a moderate and transient increase in EMG activity during REM sleep.**

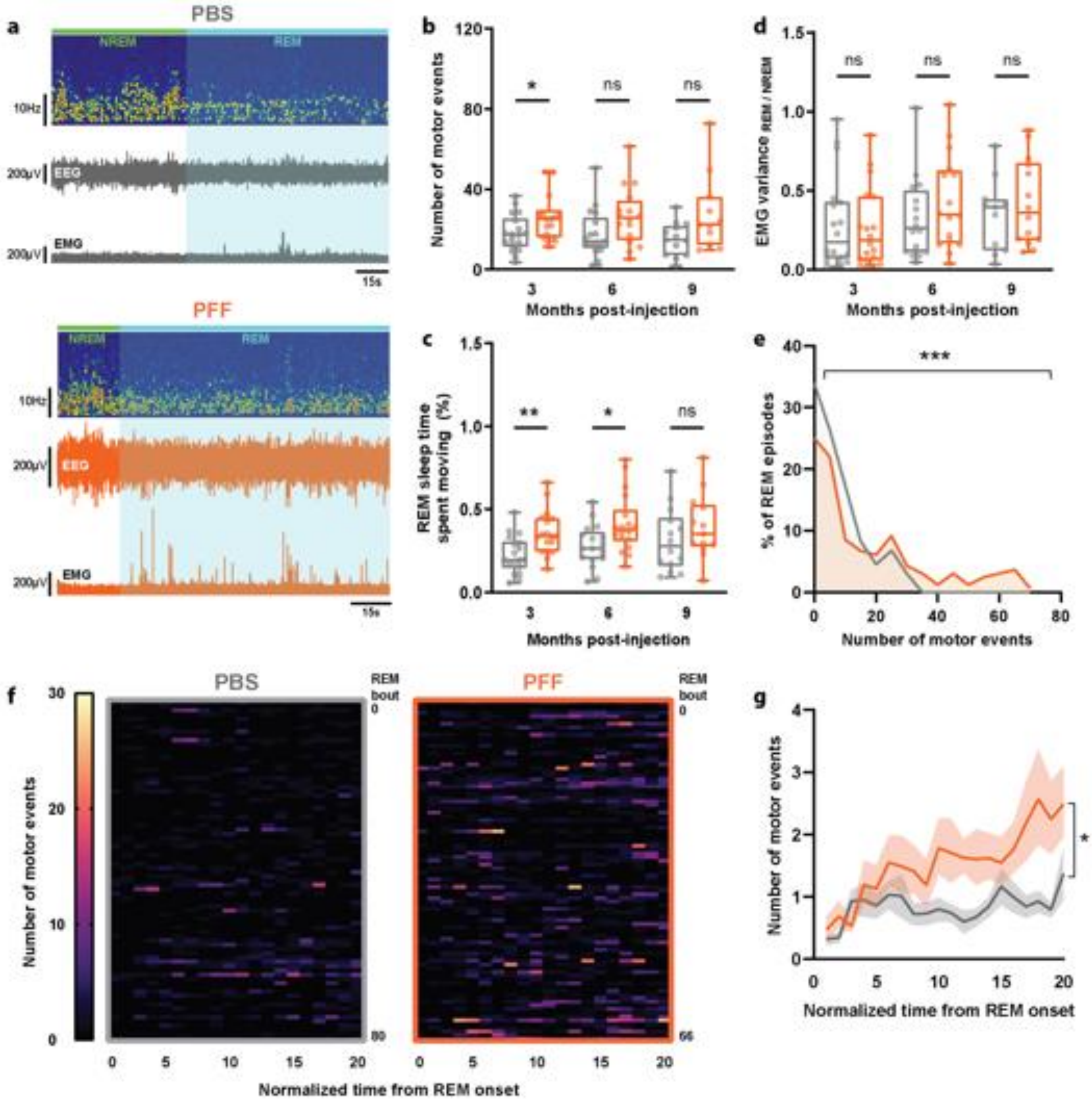


(**a**) Electrophysiological recording from mouse that received unilateral injection of PFFs into the right SLD at 9 mpi (orange) and age-matched PBS control (grey). (**b-c)** Quantitative EMG analysis revealed an increase in REM sleep movements at 3 mpi (**b**) (**P* < 0.05, *n* = 18), which was accompanied by an increase in the time that mice spent moving during REM sleep at 3 and 6mpi (**c**) (*p<0.05, **p<0.01 n=18). (**d**) The REM:NREM ratio of EMG variance did not differ between PFF-injected mice compared to age-matched controls (*P* > 0.05, *n* = 18). (e) The frequency distribution of REM sleep episodes was skewed towards episodes with more movements at 6 mpi (****P* < 0.001, *n* = 133-164 episodes). (**f**) Heatmaps show the number of movements along the standardized duration of each REM sleep episode from PBS (left) and PFF-injected mice (right) at 6 mpi. (**g**) Group data comparing the temporal profile of movements within individual REM sleep episodes from PFF-injected mice (orange line) and age-matched PBS controls (grey line) (**P* < 0.05, *n* = 18).

**Supplementary Fig. 4. PFF injection caused no changes in muscle activity during wakefulness or NREM sleep.**

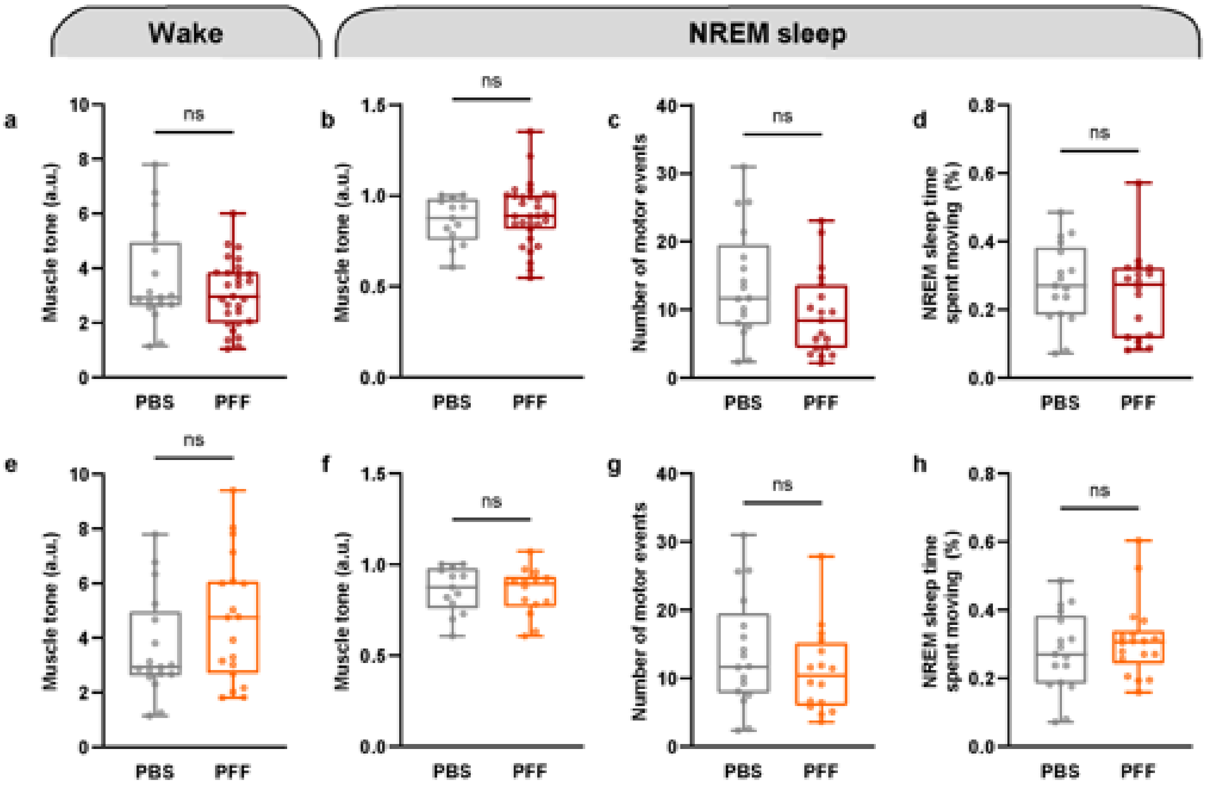


(**a-d**) Mice bilaterally inoculated with PFFs in the SLD did not show differences in muscle tone during wakefulness (**a**) or NREM sleep (**b**). In contrast with results observed during REM sleep, both the number (**c**) and the time (**d**) occupied by movements during NREM sleep were unaffected by PFF injection until 9 mpi ($P > 0.05$, $n = 18\text{-}22$). (**e-h**) Unilaterally injected mice also did not show any significant differences in EMG activity during wakefulness or NREM sleep ($P > 0.05$, $n = 18$).

**Supplementary Fig. 5. Male and female PFF-injected mice did not exhibit clear sex differences in the expression of RBD-like behaviors.**

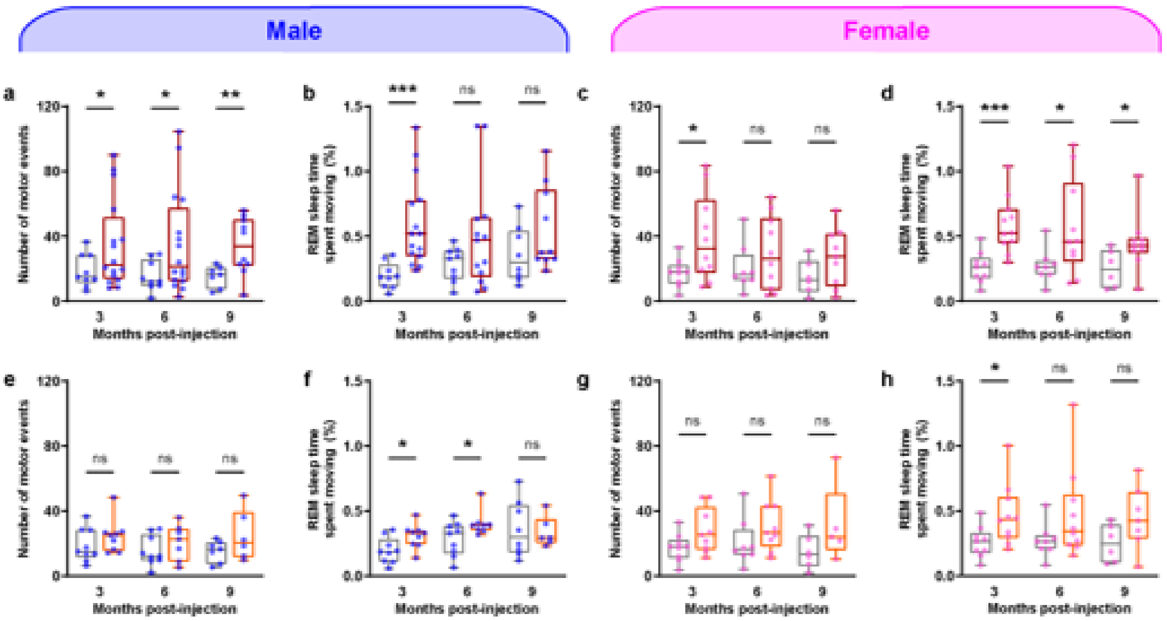


**(a-b)** The number of movements (**a**) and the time that mice spend moving during REM sleep (**b**) is shown for male mice with bilateral PFF inoculations of the SLD (*$P < 0.05$, **$P < 0.01$, ***$P < 0.001$, $n$ = 9-16). **(c-d)** The same parameters are shown for female mice that received bilateral inoculations (*$P < 0.05$, ***$P < 0.001$, $n$ = 8-10). **(e-h)** EMG data is shown for male and female mice that received unilateral PFF injections (*$P < 0.05$, $n$ = 9 males and 9 females). Note that no single EMG parameter appears to be driven by either male or female mice among either bilateral or unilaterally injected mice.

**Supplementary Fig. 6. Sex differences in gait dysfunction.**

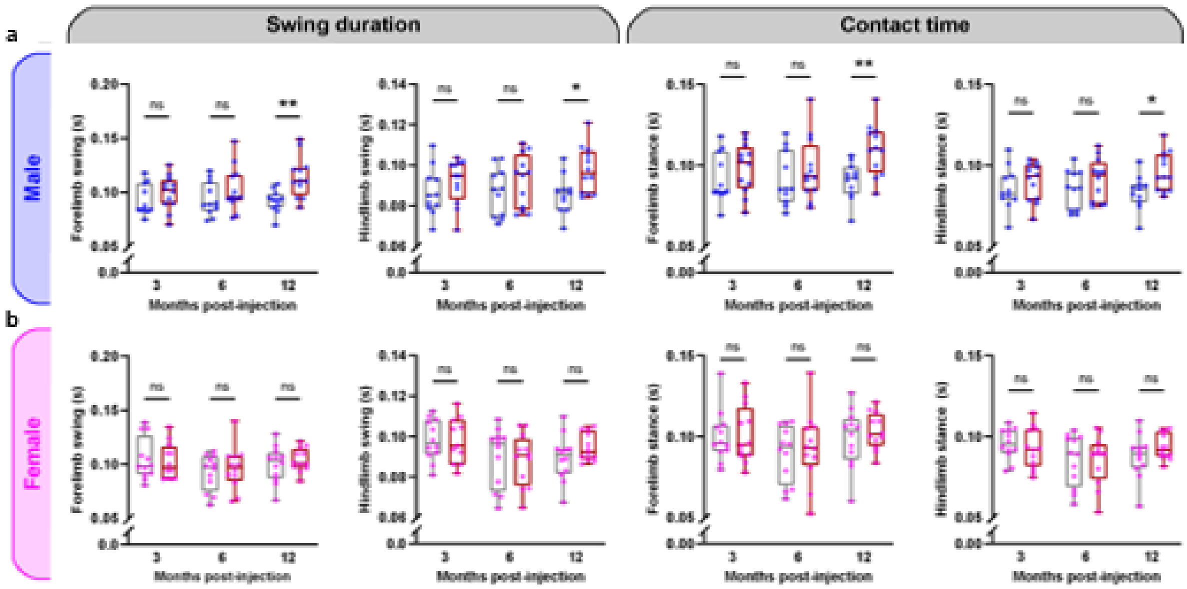


(**a**) Male mice showed significant increases in forelimb and hindlimb swing duration and contact time at 12 mpi (*$P < 0.05$, **$P < 0.01$, $n$ = 11-12). (**b**) Female mice did not differ significantly from controls in gait parameters ($n$ = 12).

**Supplementary Fig. 7. PFF-injected mice do not display severe motor or cognitive deficits**.

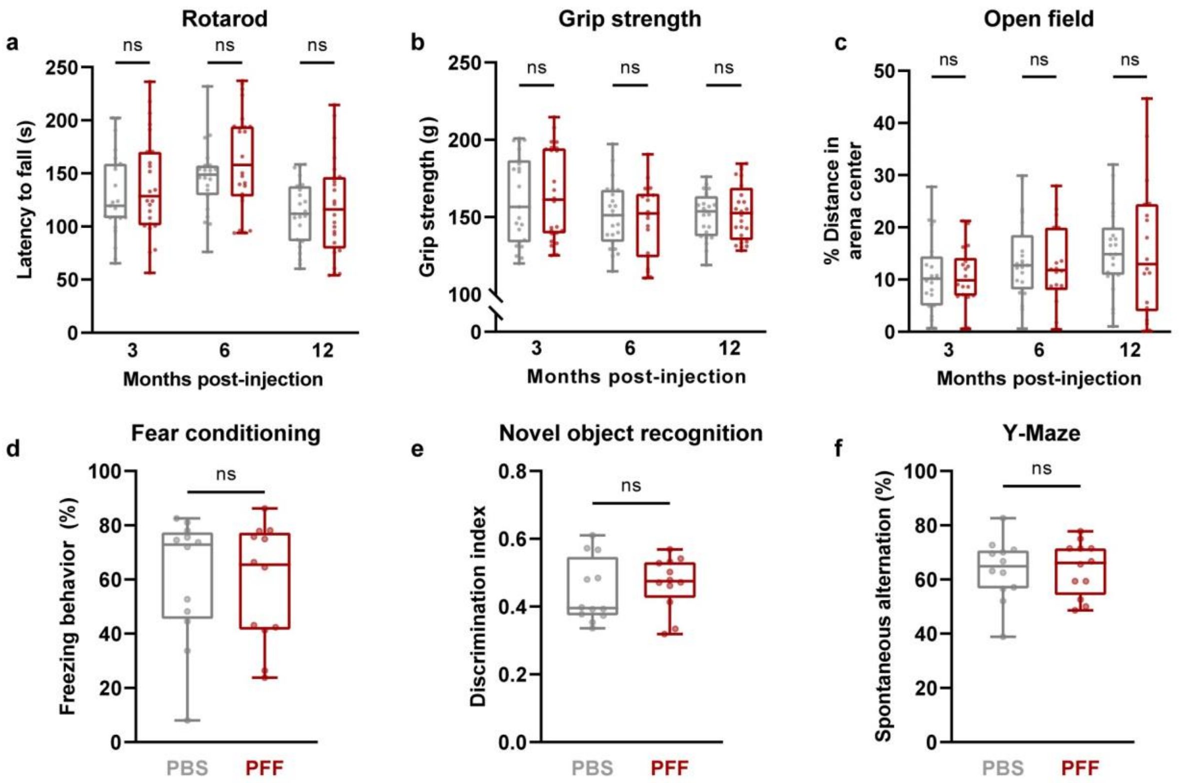


(**a**) PFF-injected mice performed similarly to age-matched controls on the accelerating rotarod, showing similar fall latencies as controls between 3 and 12mpi ($P$ = 0.2669, $n$ = 24). (**b**) Grip strength testing did not reveal differences in total grip strength compared to controls ($P$ = 0.2733, $n$ = 24). (**c**) PFF-injected mice did not display deficits on open field testing compared to age-matched controls. (**d-f**) PFF-injected mice were also tested on various cognitive tasks and do not differ from controls in freezing behaviors in fear conditioning (**d**) ($P$ = 0.8318, $n$ = 12), preference for novel objects (**e**) (p=0.4450, n=12), or spontaneous alternation behaviors on the Y-maze test (**f**) (p=0.9903, n=12) at 12 mpi.

**Supplementary Fig. 8. Association between neuronal pathological burden and the degree of RBD-like behavior**

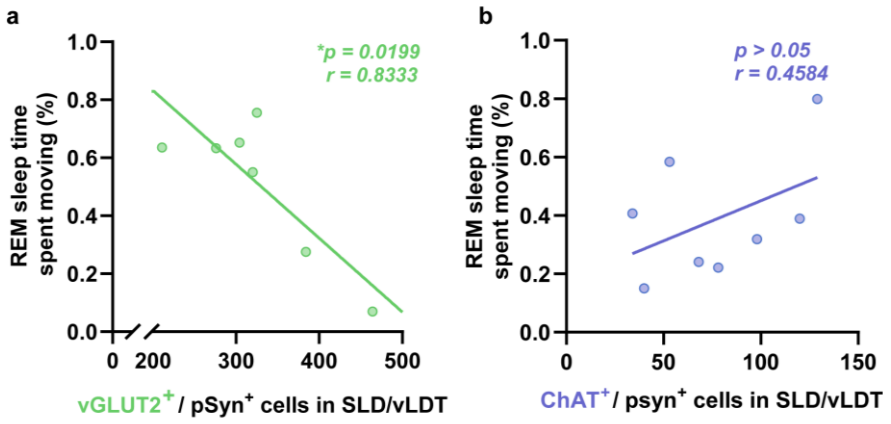


**(a)** Linear regression analysis reveals a negative correlation between the number of glutamatergic SLD/vLDT neurons and RBD-like behaviors among PFF-injected mice (*$P$ < 0.05, $n$ = 7). Mice with the most fewest remaining SLD/vLDT glutmate neurons had the highest levels of muscle activity during REM sleep. (**b**) No significant correlation is observed between the number of cholinergic SLD/vLDT neurons and RBD-like behaviors ($P$ > 0.05, $n$ = 8).

**Supplementary Fig. 9. PFF injection induced reactive gliosis in the SLD/vLDT.**

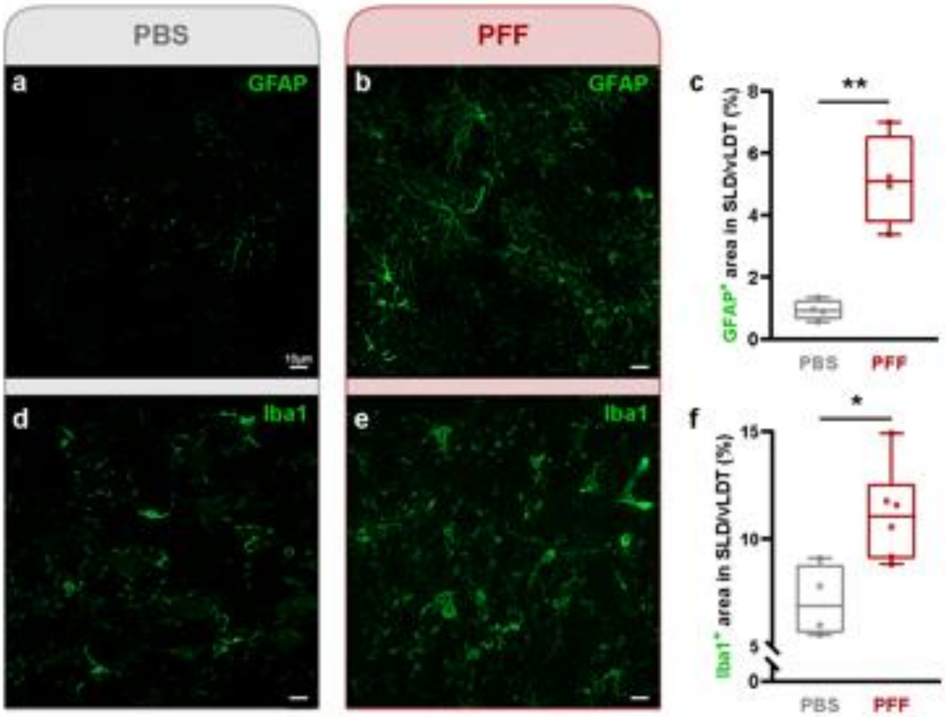


(**a**-**b**) Representative images show staining for the astrocyte marker GFAP (green) in the SLD/vLDT of a PBS-injected mouse (**a**) compared to a PFF-injected mouse (**b**) at 9 mpi. The SLD/vLDT of PFF-injected mice display greater GFAP expression compared to controls. (**c**) Quantification of histological data reveals that PFF-injected mice exhibited an increase in the GFAP-immunoreactive area within the SLD/vLDT compared to age-matched controls (*$P < 0.01$, $n = 4$). (**d**-**e**) Example micrographs show staining for the microglia marker Iba1P (green) in the SLD/vLDT of a mouse injected with PBS (**d**) or PFFs (**e**) at 9 mpi. (**f**) PFF-injected mice had a larger area of immunoreactive Iba1 within the SLD/vLDT compared to controls at 9 mpi (*$P < 0.05$, $n = 4$-$6$).

**Supplementary Fig. 10. Experiment schematic.**

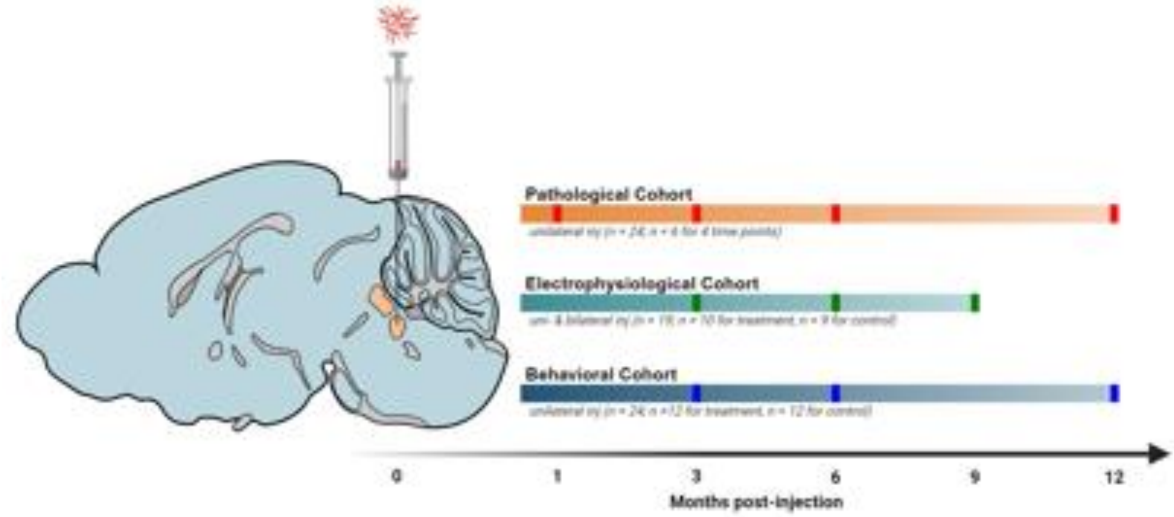


Mice were injected with αSyn in the SLD at 3 months of age. The mice were then aged 1, 3, 6, 9, and 12 mpi, and assessed for histopathology, electrophysiology, and behavior at pre-determined time points.

**Supplementary Fig. 11. Pipeline for automatic quantification of p-αSyn pathology and generation of heatmap.**

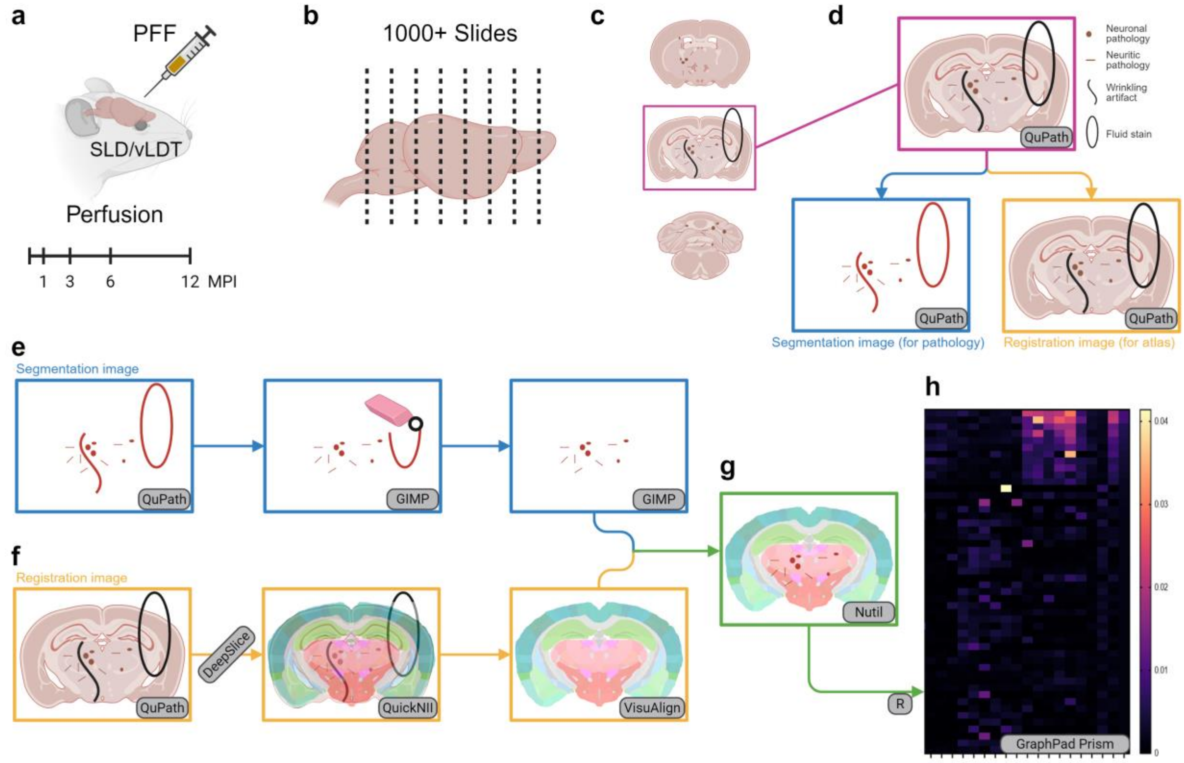


(**a**) 27 WT mice were injected at the SLD region of the brain, 24 were injected with PFF and 3 were injected with PBS for negative control. 6 PFF-injected Mice were sacrificed at 1, 3, 6, and 12 mpi, with the 3 control mice also sacrificed at the 12 mpi. (**B)** Brains were then embedded within paraffin blocks after perfusion, then subsequently cut at 6μm increments on a microtome. The whole brain was utilized, generating approximately 1000 physical slides per brain. (**c**) Of the 1000, 80 representative slides were chosen per brain, stained, and scanned. (**d**) Scanned whole-slide images were imported into QuPath to generate two subsets of images in PNG format compatible with a workflow modified from QUINT: "registration" images (low-resolution full color PNGs for Allen Brain Atlas registration) and "segmentation" images (full-resolution binary PNGs containing only pathology-positive pixels). (e) Segmentation images were manually filtered for artifacts in GIMP, which primarily consisted of non-specific edge staining. (**f**) Registration images were given to DeepSlice for a first pass at region localization, then manually adjusted for increased accuracy in QuickNII and VisuAlign. (**g**) Image sets were fed into the NUTIL quantifier function to generate a pathology pixel count for each brain region. An R script was created to compile the data for each brain across the study, calculate load metrics (pathologic area/total region area,) and to filter out irrelevant regions. Where noted, sub-regions defined by the ABA were combined into their parent features. (**h**) Resultant data was used to generate heatmaps in GraphPad Prism (version 10).
